# Highly efficient anogenital transmission of clade Ia mpox virus associated with increased shedding

**DOI:** 10.1101/2025.08.14.669880

**Authors:** Franziska K. Kaiser, Reshma K. Mukesh, Shane Gallogly, Jonathan Schulz, Sarah van Tol, Natalie McCarthy, Missiani Ochwoto, Kwe Claude Yinda, Atsushi Okumura, Lara Myers, Aaron Carmody, Brian J. Smith, Greg Saturday, Carl Shaia, Kyle Rosenke, Julia R. Port, Vincent J. Munster

**Affiliations:** Laboratory of Virology, Division of Intramural Research, National Institute of Allergy and Infectious Diseases, National Institutes of Health, Rocky Mountain Laboratories, Hamilton, MT, USA; Research and Technologies Branch, Division of Intramural Research, National Institute of Allergy and Infectious Diseases, National Institutes of Health, Hamilton, MT, USA; Rocky Mountain Veterinary Branch, Division of Intramural Research, National Institute of Allergy and Infectious Diseases, National Institutes of Health, Rocky Mountain Laboratories, Hamilton, MT, USA; Laboratory of Transmission Immunology, Helmholtz Centre for Infection Research (HZI), Braunschweig, Germany

## Abstract

The transmission pattern of mpox has shifted from sporadic zoonotic outbreaks to sustained human-to-human spread. Epidemiological data indicate sexual contact as a crucial driver for efficient transmission and the associated devastating mpox outbreaks in recent years. However, our understanding of exact driving factors and transmission determinants is still limited. Here, we investigated MPXV clade Ia virus pathogenicity, shedding kinetics, and transmission potential in a prairie dog model (*Cynomys ludovicianus*). All tested mucosal inoculation routes (penile/preputial, vaginal, rectal, intranasal) resulted in a productive, systemic infection. Inoculation via urogenital routes generated the highest virus shedding and most severe clinical disease. A simulated sexual contact transmission resulted in 100% transmission efficiency with high virus shedding in sentinels on day 1, even before the onset of clinical signs. Our findings provide critical insights into mpox transmission, emphasizing the role of anogenital mucosal surfaces in facilitating rapid spread. These results advocate for a stronger focus on mucosal infection when evaluating countermeasures.

## Introduction

Since 2022, mpox has caused an increasing global Public Health burden^1,2^. The etiological agent of mpox, monkeypox virus (MPXV), belongs to the same genus as smallpox, *Orthopoxvirus*, and is divided into clade I and clade II^3^. Clade I (formerly known as Congo Basin clade), is generally associated with a higher case fatality rate compared to a milder disease progression in clade II (formerly known as West African clade)^4^. Until 2022, mpox outbreaks were located mainly in west and central Africa and comprised of small case clusters after zoonotic transmission with limited human-to-human transmission after close contact^5^. This paradigm shifted in May 2022 with the rapid spread of mpox cases to previously non-endemic countries^6,7^.

In 2023, reports from the Democratic Republic of the Congo (DRC) raised concerns about a rising number of human infections with MPXV clade I^8,9^. Mpox was declared a public health emergency of international concern for the second time by the WHO on August 14^th^, 2024^10^. The current spread of clade I is mainly driven by efficient close contact transmission with different strains (clade Ia and clade Ib) simultaneously co-circulating after multiple independent zoonotic introductions^11^. Additionally, the newly emerged clade Ib has spread to central and east Africa and has caused travel-associated cases in other African countries, Europe, America, and Asia^12^. Moreover, phylogenetic analysis has shown increased APOBEC3 editing in clade Ia and Ib strains, a mutation signature associated with continuous circulation within the human population^11,13,14^.

Epidemiological investigations have traced transmission events during the ongoing clade I outbreak to close, mainly physical contact between household members, relatives, colleagues, or sexual networks^15^. However, many questions about conserved and strain-specific transmission routes, virus shedding kinetics, and modes of human-to-human spread remain unanswered, as epidemiological data is currently still limited^16^.

Here, we investigated mpox clade Ia virus shedding kinetics, pathogenicity, and transmissibility in dependence on different mucosal inoculation sites in black-tailed prairie dogs (*Cynomys ludovicianus*). Mpox infections in prairie dog model resembles human clinical disease, including short febrile prodromal phase followed by generalized pox lesions^17,18^. Other available rodent models such as castaneous mice, which have an impaired innate immune response, succumb to the infection without displaying a course of disease progression that is comparable to human patients^19,20^. Prairie dogs have been successfully used for translational studies to test the efficacy of vaccinations and antiviral treatments against mpox challenge^21-23^. Furthermore, mpox transmission has also been described between prairie dogs in classical experimental settings^24^, as well as during the 2003 outbreak in US pet prairie dogs^25^.

This study demonstrates the efficiency of anogenital infections, as well as rapid onset and high titer virus shedding after simulated sexual contact transmission.

## Materials and Methods

### Ethics statement and biosafety

Animal experiments were approved by the Institutional Animals Care and Use Committee and performed in an AAALAC accredited animal holding facility. Prairie dogs were housed in climate-controlled rooms with fixed 12-hour light-dark cycles. The animals were monitored at least once daily.

This project was reviewed and approved by the NIH RML Institutional Biosafety Committee (IBC) and classified as BSL3. In addition, the NIH Dual Use Research of Concern-Institutional Review Entity (DURC-IRE) concluded that the project did not meet the US Government definition of DURC or Enhanced Potential Pandemic Pathogens (ePPP). All experiments were performed under maximum containment conditions following Institutional Biosafety Committee (IBC) approved standard operating and inactivation procedures.

### Viruses and cells

Vero E6 cells (provided by Ralph Baric, University of North Carolina) were maintained in Dulbecco’s Modified Eagle Medium (DMEM) supplemented with 10% fetal bovine serum (FBS), 1□mM L-glutamine, 50□U□ml^−1^ penicillin, and 50□µg□ml^−1^ streptomycin. For growing virus stocks and plaque assays, cells were cultured in DMEM with 2% FBS 1□mM L-glutamine, 50□U□ml^−1^ penicillin, and 50□µg□ml^−1^ streptomycin (DMEM2). MPXV Zaire 79 (V79-I-005) was obtained from BEI resources (https://www.beiresources.org/Catalog/animalViruses/NR-2324.aspx). The virus was sequenced to confirm the sequence and exclude contamination.

### Exposure route comparison in prairie dogs

Black-tailed prairie dogs (*Cynomys ludovicianus*) were randomly assigned to four groups (*n=4*, mixed sex for nasal and rectal exposure). All groups were inoculated with 2.5×10^4^ PFU MPXV Zaire-79 via different mucosal routes (penile/preputial, vaginal, intranasal, rectal). Prior to inoculation, a dry sampling swab (Polyester tipped applicator, 25-800 1PD, Puritan, ME, USA) was used to remove mucus and cause microabrasions of the mucosa. The inoculation volume was 200 µl for the rectal, and 20 µl for the nasal, penile/preputial and vaginal group. The inoculum was carefully pipetted into the respective orifice. Control animals underwent the same procedures and were rectally mock inoculated with 200µl DMEM. All animals were handled under isoflurane anesthesia.

### Experimental design transmission study

To determine the transmission dynamics, four animals per group (Donors) was inoculated rectally and nasally with 2.5×104 PFU MPXV Zaire-79 as described above. We simulated mucosal contact transmission by using a swab to transfer virus from a donor to a naïve sentinel animal (Figure 5A). 11 days post inoculation, a swab was taken from the rectum of a rectally infected donor animal and was immediately transferred to the rectum of a sentinel (4 transmission pairs). Prior to the transfer, the rectum of the sentinel animal was cleaned with a dry swab to remove mucus and cause microabrasions. The transfer swab was inserted into the sentinel’s rectum and left in place for 5 min. Similarly, a nose-to-nose swab transfer was conducted. Nose swabs from 4 intranasally inoculated donors were collected on day 11 and immediately transferred to the nose of 4 sentinel animals. Again, sentinels were prepared for the transfer by removing residues or mucus from their nostrils before placing the swab in the outer part of their nostrils for 5 min while the animals were anesthetized. The experiment included 4 transmission pairs per transfer route. All animals were housed individually during the study. Virus titer of transfer swabs was determined by plaque assay.

### MPXV quantitative real-time PCR

For nucleic acid extraction from animal tissues, organ material was weighted and homogenized in RLT buffer. Swab samples were vortexed in DMEM2 and 140µl were used for extractions with the QIAamp Viral RNA Kit (Qiagen) kit according to the manufacturer’s description. MPXV DNA and RNA were detected using a MPXV qRT-PCR assay for the detection of the G2R gene coding sequence^26^. In brief, isolated nucleic acids were run using a TaqMan Fast Virus One-Step Mastermix (Applied Biosystems) with 15µl reaction mix and 5µl extracted nucleic acids. The reaction mix contained 10µM primers each and 10µM probe. Forward primer 5’-GGAAAATGTAAAGACAACGAATACAG-3’, reverse primer 5’-GCTATCACATAATCTGGAAGCGTA-3’ and probe 5’-FAM-AAGCCGTAATCTATGTTGTCTATCGTGTCC-3’-BHQ1. Primers and probes were sourced from Integrated DNA Technologies (Coralville, Iowa, USA). qRT-PCR reactions were performed on QuantStudio 3 thermocycler. MPXV standards with known copy numbers were used to calculate the standard curve.

### Infectious virus quantification

Virus titer in swab samples was determined via endpoint plaque assay with ten-fold serial dilutions in DMEM with 2% FBS. 400µl of each dilution was added into a well of a 48-well plate, coated with confluent Vero E6 cells. After four days at 37°C with 5% CO_2_ plates were fixated with 10% formaldehyde for 10 min. Subsequently, the formaldehyde was removed, and plates were stained for additional 10 min with a 1% crystal violet solution. After rinsing and drying, plaques were counted and virus titers calculated.

### Histopathology

Tissues were placed in cassettes and fixed for a minimum of 7 days in 10% neutral buffered formalin. Subsequently, tissues were processed using a graded series of ethanol, xylene, and PureAffin, with a Sakura VIP-6 Tissue Tek, on a 12-hour automated schedule. Embedded tissues are sectioned at 5µm and dried overnight at 42°C prior to staining. MPXV antigen was detected by immunohistochemistry (IHC). As a primary antibody GeneTex vaccinia virus antibody (#GTX36578) was applied. Vector Laboratories ImmPRESS-VR horse anti-rabbit IgG polymer (# MP-6401) was used as a secondary antibody. The tissues were stained using the Discovery Ultra automated stainer (Ventana Medical Systems) with a Roche Tissue Diagnostics Discovery purple kit (#760-229). Tissues were evaluated for histological lesions.

### Flow cytometry

Prairie dog spleens were harvested and brought to single cell suspension by gentle mechanical homogenization through a 100-µm nylon filter, followed by red blood cell lysis with a 5 min treatment at RT in ACK Lysing Buffer (Gibco). Cells were washed in RPMI-1640 media. Then, 1-2×10^6^ splenocytes were plated in 96-well U-bottom plates. Prior to surface staining, cells were stained for 20 min at RT with the Fixable Blue Dead Cell Stain Kit (L34962, Invitrogen) to identify viable cells and then quenched/washed with a 2% FBS-PBS wash buffer. The antibodies used for direct ex vivo surface staining were BV605-anti-CD4 (L200; 562843, BD Biosciences), APC-ef780-anti-HLA-DR (LN3; 47-9956-42, Invitrogen), BUV395-anti-CD21 (B-ly4; 740288 BD Biosciences), FITC-Goat anti-Guinea Pig IgG (H/L) (polyclonal; AHP863F, BioRad) and Pacific Blue-anti-CD14 (Tuk4; MHCD1428, Life Technologies). Samples were stained at 4°C for 30 min, then washed in 2% FBS-PBS wash buffer and fixed overnight using the eBioscienceTM Foxp3/Transcription Factor Staining Buffer set (00-5523-00, ThermoFisher). Samples were then permeabilized using this kit and stained for the intracellular expression of FITC-anti-CD3 (CD3-12; MCA1477F, BioRad) or AF647-anti-CD3 (CD3-12; MCA1477A647, BioRad), PE-anti-CD79a (HM47; 555935, BD Biosciences), PerCP/Cy5.5-anti-Ki-67 (B56; 561284, BD Biosciences) or R718-anti-Ki-67 (B56; 566963, BD Biosciences) and PerCP/Cy5.5-anti-CD68 (FA-11; 137010, BioLegend) for 30 min at 4C. Samples were then washed using perm buffer then fixed again overnight in 2% paraformaldehyde (PFA). Samples were then centrifuged, transferred to fresh 2% PFA and then centrifuged and transferred to 2% FBS-PBS wash buffer. Data was analyzed using a FACSymphonyTMA5 (BD Biosciences) and FlowJo software (version 10.9.0; BD Biosciences).

### Luminex MagPlex serology

Recombinant antigens (Sino Biological) were individually coupled to xMap MagPlex-C microspheres (Luminex) at 10 ug per bead region using the BioPlex Amine Coupling Kit (Bio Rad) according to the manufacturer’s instructions. Positive control microspheres were prepared by coupling 10 ug Protein G (Thermo Fisher) to one region, and negative control microspheres were coupled to 10 ug human serum albumin (Sigma Aldrich) to one bead region. All regions were combined and pipetted into a 96 well plate at an estimated 500 beads per region per well. Serum samples were applied at a dilution of 1:100 in staining buffer (Bio Rad), and bound antibody was detected with a mixture of biotinylated protein G and protein A (Thermo Fisher, 2 ug/mL) diluted in staining buffer. Streptavidin-PE (Southern Biotech) was applied at 2 ug/mL diluted in staining buffer, and the beads were resuspended in PBS-Tween prior to reading the plate. Analysis was done using the MAGPIX System and Luminex xPONENT for MAGPIX software. Results were reported as median fluorescence intensity (MFI), analyzing a minimum of 50 microspheres per region.

### Statistical analysis

Data visualization and statistical tests were performed using GraphPad Prism version 10.2.0. Exact tests for significances and p-values are indicated where appropriate.

## Results

### Mucosal MPXV infection causes systemic virus dissemination and severe disease phenotype

To better understand the influence of the initial MPXV entry site on pathogenicity and disease progression, we first compared four mucosal inoculation routes tailored to model human sex-associated contact. Black-tailed prairie dogs (*Cynomys ludovicianus*) were randomly assigned to the four groups and inoculated with 2.5×10^4^ PFU MPXV clade Ia via the intranasal, vaginal, penile, or rectal route (Figure 1A). Animals were euthanized on day 11 post inoculation upon displaying signs of peak disease. Clinical signs in infected animals included redness and swelling of inoculation site for anogenital inoculation routes. Initial clinical signs were observed 5-9 dpi. Generalized skin lesions were observed in 13 of 16 animals (81.25%), starting on day 9-11. Body temperature peaked on day 7 with significantly higher temperatures for the vaginal and penile inoculation compared to the control animals (Figure 1C). First animals from vaginal, penile, and rectal groups reached humane endpoint criteria on day 11, which was defined as study endpoint. Weight loss during the study was minimal and did not serve as a reliable indicator for disease severity (Supplemental Figure 1).

**Figure 1:**
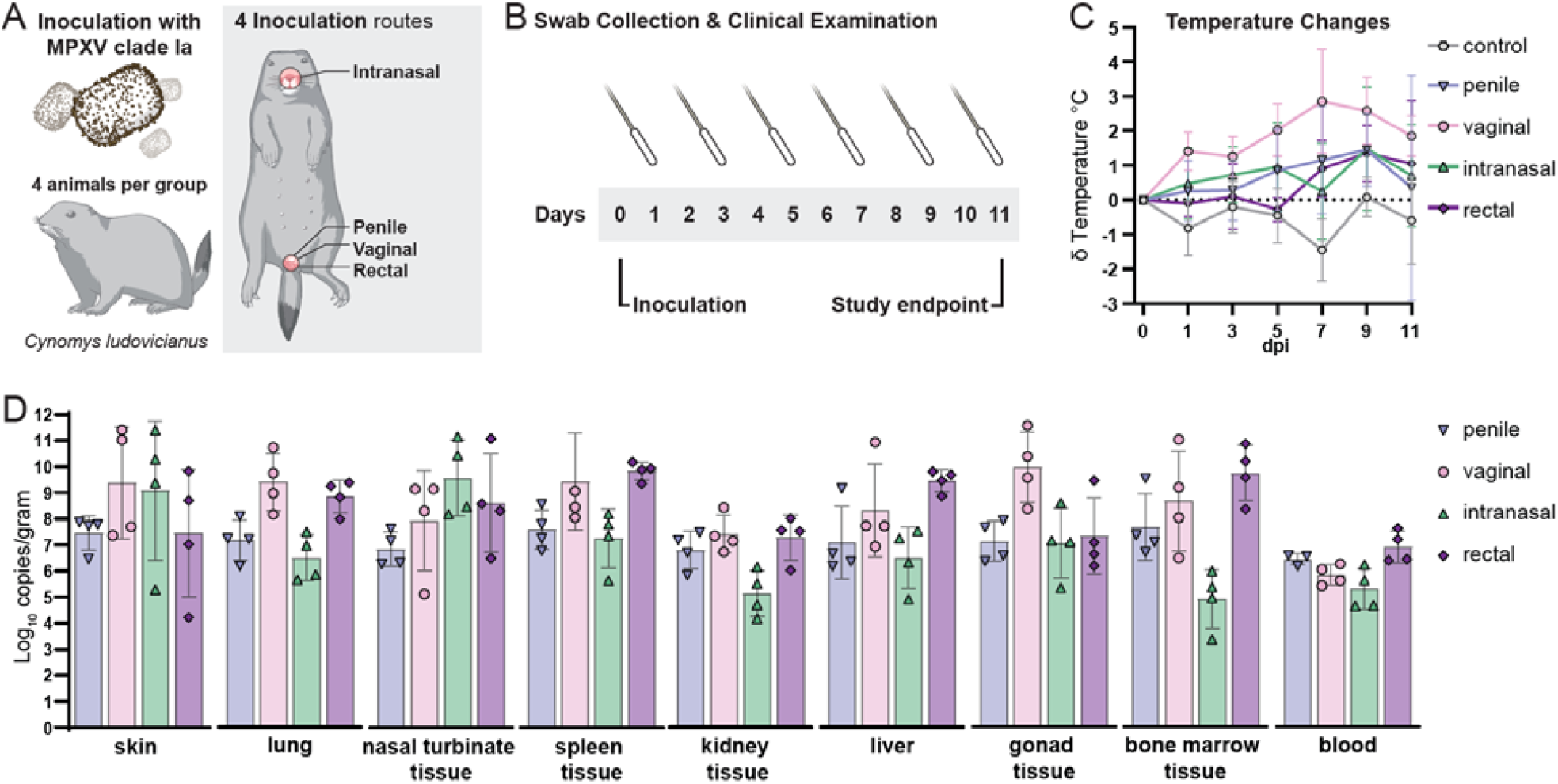
Experimental design of exposure route comparison, relative body temperature changes over time, and MPXV organ dissemination. (A) Black-tailed prairie dogs (*Cynomys ludovicianus*), 4 per group, were inoculated with 2.5×10^4^ PFU MPXV Zaire 79 (V79-I-005). Animals were inoculated via the penile, vaginal, rectal, or nasal routes. (B) On day 1, 3, 5, 7, 9, and 11, clinical exams were performed., Body weight, temperature, as well as oropharyngeal, nasal, urogenital, and rectal swab samples were taken. (C) Body temperature was compared to rectally mock-inoculated control animals (mean; error bars show standard deviation). (D) Organ dissemination was assessed by quantifying virus genetic copy numbers at study endpoint 11 days post inoculation using qRT-PCR. The bars represent mean values and standard deviation. Individual data points are displayed as symbols.

Next, we determined the organ dissemination at the study endpoint (11dpi). MPXV loads in organ tissue were quantified by qRT-PCR for lung, nasal turbinates, liver, spleen, kidney, skin, gonads, and bone marrow. All organs tested positive for MPXV genomic material in all animals independent of the inoculation route. This demonstrates systemic spread and complete dissemination after mucosal infection. The highest virus loads were detected in parenchymal organs (liver, spleen, kidney) and bone marrow for the rectally inoculated groups. The lowest viral loads were observed in the intranasally challenged animals, except for nasal turbinates and skin samples. Skin samples were taken from the animal’s chin, where skin lesions first occurred and showed the highest variability in viral loads (Figure 1D).

Taken together, prairie dogs were equally susceptible for clade Ia MPXV regardless of inoculation route, as demonstrated by a marked systemic dissemination and viremia. However, anogenital routes of inoculation resulted in a more severe disease characterized by highest viral loads in swab samples and organ tissue.

### Distinct MPXV pathological lesions throughout several organ systems

Next, we investigated whether the different exposure routes would result in distinct pathological manifestations. Post-mortem examinations revealed moderate to severe localized inflammation with purulent discharge at the inoculation site for all experimental groups. After rectal and genital inoculation, lymphoid tissues, including genital- and urinary tract-associated structures were mild to markedly enlarged. This caused 2 of out 4 vaginally challenged animals difficulties passing urine.

Histopathologically, the respective inoculation sites were characterized by inflammation, ulceration and necrosis of epithelial, sub-epithelial and deeper tissues including muscle (myonecrosis) and adipose tissue (steatonecrosis) (Figure 2). Inflammation in affected and adjacent tissues was severe and consisted of viable and degenerate neutrophils, macrophages and fewer lymphocytes. Glandular epithelial cells demonstrated characteristic ballooning degeneration (intracellular edema) and eosinophilic intracytoplasmic inclusion bodies. Immunohistochemistry staining demonstrated concomitant presence of viral antigen in affected areas and adjacent non-affected tissues. All experimental groups presented with varying degrees of inflammation and necrosis, ranging from minimal to marked in systemic organs including the spleen, liver, lung, and bone marrow.

**Figure 2:**
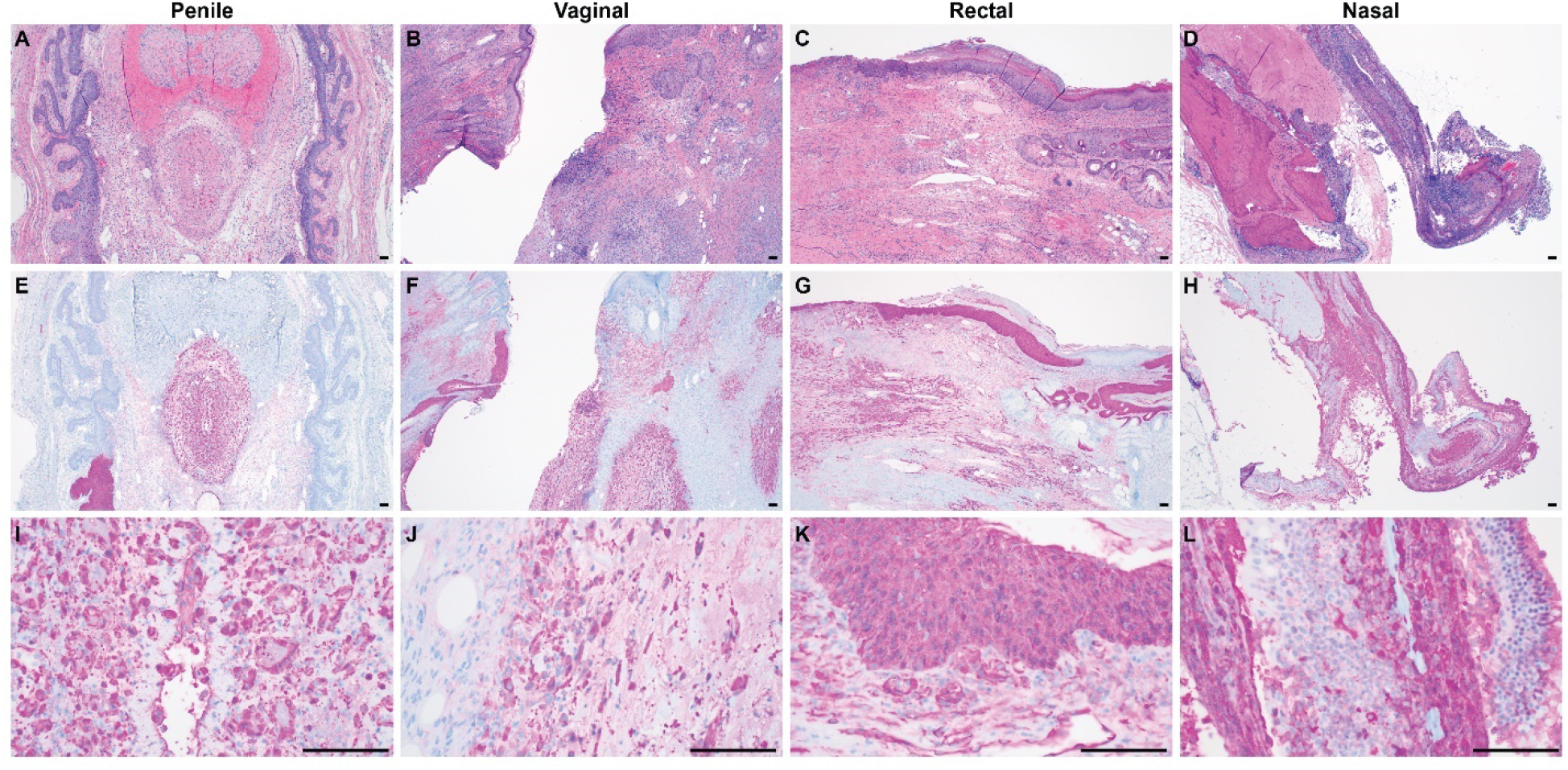
Histology revealed severe inflammation and necrosis of the mucosa and underlying tissues at the inoculation site following MPXV challenge. (A,E,I) Penis and prepuce after penile inoculation displayed ulceration of the penile urethra with circumferential inflammation, necrosis and edema extending into the submucosa and tunica albuginea separating the urethra from the corpus spongiosa and corpus cavernosa. Immunohistochemistry (IHC) images of glandular epithelial cells, fibroblasts and inflammatory cells demonstrated MPXV antigen presence. (B,F,J) Vaginal inoculation resulted in ulceration of the vaginal mucosa with inflammation and edema of the submucosa extending into surrounding tissues. Remaining epithelial cells, fibroblasts and inflammatory cells expressed positive signal in IHC staining, as does a portion of intact, hyperplastic epithelium. (C,G,K) Ulceration of the rectal mucosa with subjacent inflammation fibroplasia and edema of the submucosa and deeper tissues and adjacent epithelial hyperplasia were visible following rectal inoculation. Epithelia, reactive fibroblasts, endothelial cells and inflammatory cells show MPXV antigen presence. (D,H,L) Intranasal inoculation resulted in ulceration of a portion of the olfactory epithelium with subjacent inflammation and fibroplasia. MPXV was detected in submucosal glandular cells, endothelial cells and fibroblasts. Panels A-D are representative images of hematoxylin and eosin-stained mucosal tissues from the inoculation site (40x magnification), E-H (40x magnification) and I-L (400x magnification) are immunohistochemistry (IHC) images using a polyclonal IgG vaccinia virus antibody. Scale bars indicate 100µm.

Histopathology demonstrated moderate to severe localized inflammation around the inoculation site along with systemic virus dissemination independent of the inoculation route.

### Anogenital entry routes are associated with increased virus shedding

Next, we assessed infectious virus shedding kinetics. Swab samples (oropharyngeal, nasal, urogenital, and rectal) were taken on days 1,3,5,7,9, and 11 (Figure 1B). All inoculation routes resulted in substantial virus shedding at the corresponding mucosal sites as detected in oropharyngeal, nasal, urogenital, and rectal swabs throughout the study period (Figure 3). Virus shedding from the inoculation sites was detected as early as day 3 in the anogenitally infected groups. In contrast, virus shedding from the inoculation site was delayed until day 5 in the nasally inoculated group. All animals inoculated via anogenital routes shed consistently high amount of virus from the respective mucosal sites throughout the study period. However, intranasally inoculated animals shed relatively lower amount of virus, and shedding was intermittent. Majority of the animals inoculated via penile and vaginal routes started shedding virus from the oral, nasal and genital sites, beyond the original inoculation site, from day 5 post-inoculation. In contrast, majority of the rectally inoculated animals began shedding virus from other sites than the original inoculation site starting on day 7 post-inoculation. Nasally inoculated animals exhibited an intermittent pattern of shedding throughout the study period. Virus shedding peaked on day 7 post inoculation for penile and vaginal infections in urogenital swabs. Highest virus titer was reached on day 7 for urogenital swabs after vaginal inoculation (mean: 7.61×10^6^ PFU/ml), followed by day 5 in urogenital swabs after vaginal inoculation (4.55×10^6^ PFU/ml). Penile inoculation resulted in 3.36×10^6^ PFU/ml on day 7 and rectal inoculation reached 3.19×10^6^ PFU/ml. The highest infectious virus titer after nasal exposure was 1.31×10^5^ PFU/ml in oropharyngeal swabs. The exact onset of infectious virus shedding compared to onset of clinical signs is displayed in Supplemental Figure 2.

**Figure 3:**
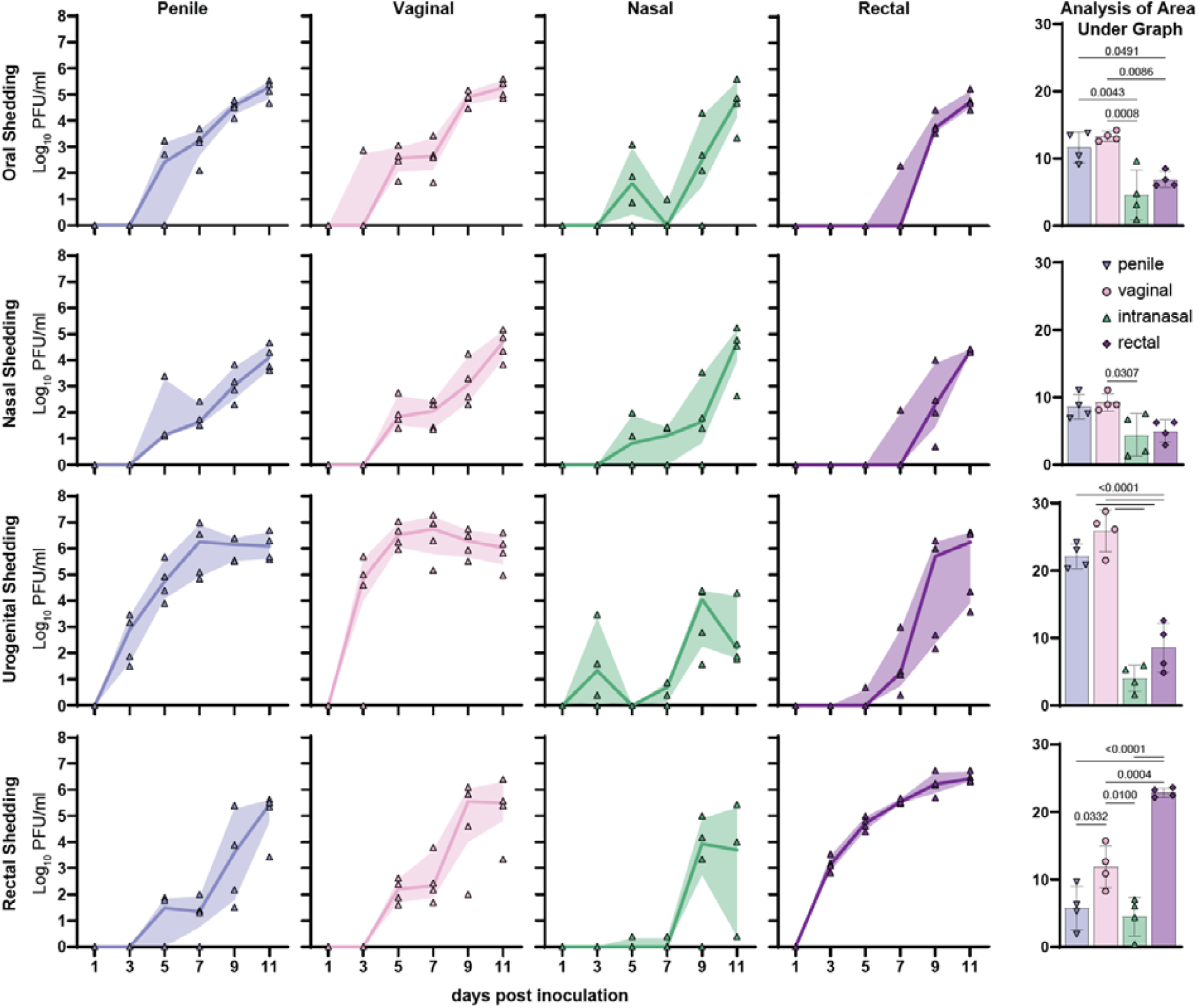
Anogenital entry routes are associated with increased virus shedding. Infectious MPXV shedding kinetics measured in Log (10) PFU/ml in oropharyngeal, nasal, urogenital and rectal swabs. Corresponding area under the curve (AUC) analyses per shedding route analyzed for statistical significance by ordinary one-way ANOVA followed by Tukey’s multiple comparison test. Infectious virus titers were determined via plaque assay. Virus shedding curves display mean. Filled area shows interquartile range. Individual data points are displayed as symbols in graph.

Area under the curve analysis for cumulative shedding demonstrated highest virus shedding for urogenital swabs after vaginal or penile inoculation (p<0.0001 compared to nasal or rectal). Similarly, rectal inoculation resulted in significantly more rectal shedding compared to all other exposure routes (p<0.004). However, intranasal inoculation did not result in higher infectious virus titers in nasal swabs compared to the other inoculation routes (mean_nasal_=4.39; mean_rectal_=5.00; mean_vaginal_=9.24; mean_penile_=8.58). Thus, shedding of infectious virus was found to be dependent on route of inoculation, with higher levels observed following genital and rectal exposure compared to nasal exposure.

### Enhanced immune cell proliferation and expansion after anogenital mpox infection

To understand the immune cell dynamics after MPXV infection, splenocytes were harvested on 11 dpi and analyzed using multi-parameter flow cytometry. An initial evaluation of commercially available antibody clones was performed on prairie dog leukocytes to assess the cross-reactivity. This screening process allowed us to achieve some basic immune cell identification and phenotyping (Supplemental Table 1 & Supplemental Figure 3). We first identified and compared the proportions of lymphocytes and myeloid cells in spleen tissues from infected and uninfected control animals based on FSC-A vs SSC-A (Figure 4A). A shift in the ratio of lymphocytes to myeloid populations was observed in infected animals compared to the uninfected control group. Control animals had an average of 91.2% lymphocytes and 7.5% myeloid cells in the spleen. In contrast, the lymphocyte-to-myeloid cell ratio shifted markedly following urogenital and rectal exposure with lymphocyte/myeloid distributions of 63.7%/30.7% in vaginally exposed animals, 63.9%/33.8% in penile-exposed animals, and 62.5%/35.4% in rectally exposed animals. The shift was less pronounced after intranasal infection, with lymphocyte and myeloid cell proportions of 73.3% and 25.2%, respectively (Figure 4B). Our results indicate a significant shift in immune cell composition following mucosal MPXV exposure, particularly after anogenital inoculation. Therefore, the magnitude in lymphocyte/myeloid ratio shifts within the immune system correlates with the magnitude of infectious virus titer, with larger shifts reflecting more infectious virus.

**Figure 4.**
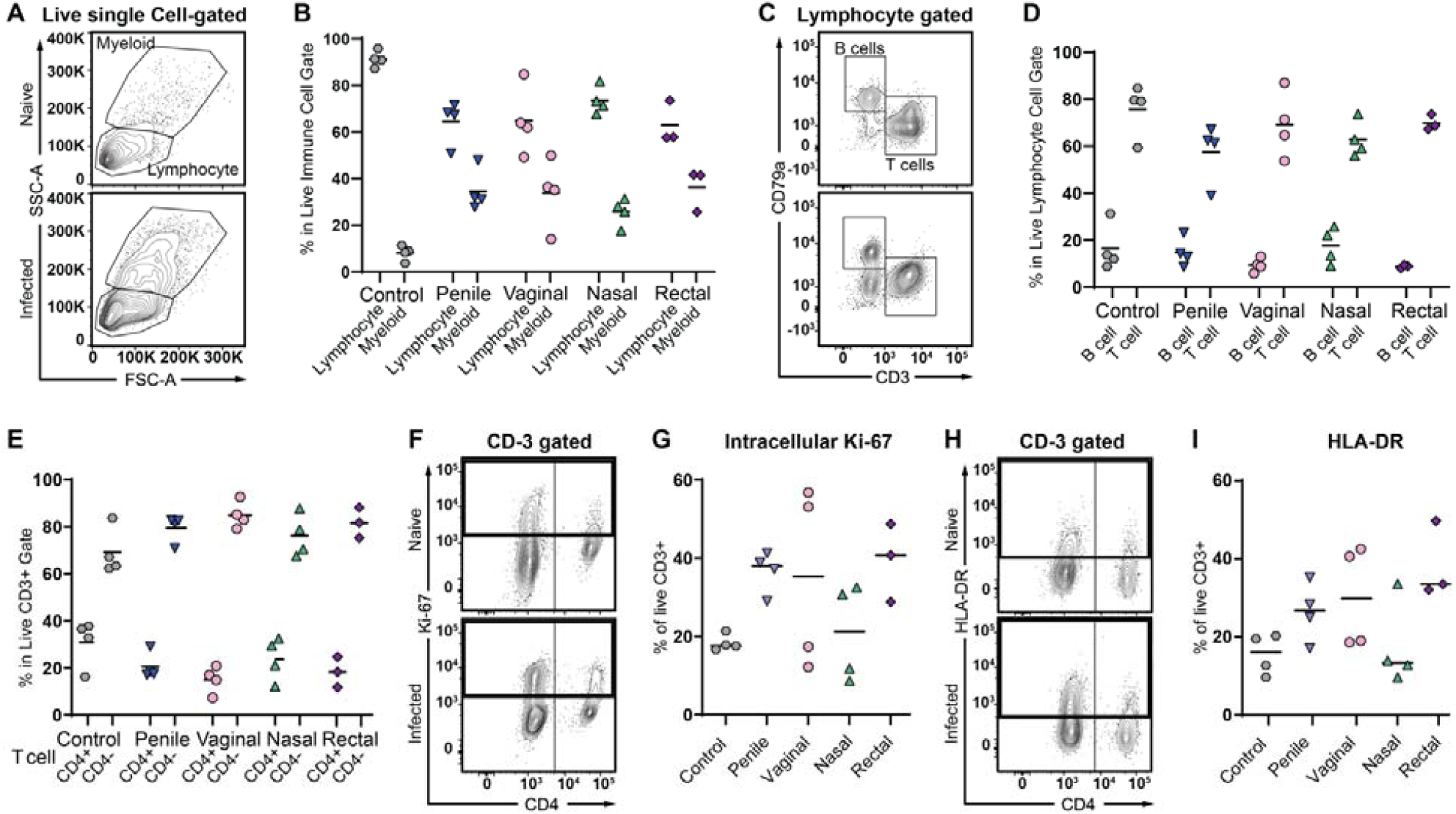
Anogenital Infection of prairie dogs with MPXV results in myeloid infiltration into the spleen with increased T cell activation. Prairie dogs were inoculated with MPXV by the penile, vaginal, intranasal and rectal route and splenocytes were analyzed on day 11 post-infection by flow cytometry. Mock-inoculated animals were used as naïve controls. (A) Representative FACS plots showing the myeloid and lymphocyte gating for naïve (upper panel) and infected (lower panel) as determined by FSC-A vs SSC-A. (B) The percentage of live myeloid-gated and lymphocyte-gated cells for the naïve control and the infected groups. (C) The lymphocyte-gated immune cells were further analyzed for intracellular expression of CD79a to identify B cells and for an intracellular CD3 epitope to identify T cells, as depicted by representative FACS plots. (D) The percentage of live B and T cells in each group. (E) The percentage of CD3^+^ T cells staining CD4^+^ and CD4^-^ (as depicted in Supplemental Figure 3C) for the naïve controls and infected groups. CD3^+^ splenic cells were further analyzed directly ex vivo for (F&G) intracellular expression of Ki-67 and (H&I) surface expression of HLA-DR. For naive controls, penile, vaginal and intranasal routes n=4. For rectal route n=3. The bar indicates the mean, and each symbol represents an individual animal.

We next analyzed the lymphocyte-gated population for T and B cells by probing intracellular markers CD3 and CD79a, respectively (Figure 4C). Compared to naive animals, no significant change in the proportion of T and B cells were observed after MPXV infection, regardless of the routes of exposure (Figure 4D). Furthermore, as we were able to identify CD4^+^ T cells from the total CD3^+^ T cell population (Supplemental Figure 3B & C), we observed a slight reduction in the proportion of CD4^+^ T cells coinciding with a slight increase in the CD4^-^ T cells for all groups post exposure (Figure 4E). In support of this observation, the intracellular proliferation marker Ki-67 (Figure 4F) and activation marker HLA-DR (Figure 4H) were highly upregulated in the CD3^-^gated CD4^-^ T cells upon MPXV exposure. As no anti-CD8 clones tested stained positive, further investigation is required to confirm the identity of these CD3^+^CD4^-^ T cells that become activated upon infection. Therefore, the total CD3^+^ T cell population was analyzed for Ki-67 and HLA-DR expression, which was strongest after anogenital exposure compared to non-infected control animals. Ki-67 expression increased from 18.3% in control animals to 36.3%, 28.3%, and 38.5% in penile, vaginal, and rectal infections, respectively (Figure 4G), while intranasal exposure was not associated with an increase (17.9%). Similarly, HLA-DR rose from 14.8% in controls to 25.6% after penile, 27.9% after vaginal, and 37.7% after rectal challenge (Figure 4I), while intranasal exposure remained at 15.5%. Luminex MagPlex serological analysis revealed only low antibody titers 11 days post inoculation (Supplemental Figure 4).

Overall, anogenital inoculation resulted in a more pronounced shift in the lymphocyte/myeloid cell ratio and increased T cell activation compared to nasal inoculation.

### Rapid onset of MPXV shedding after simulated mucosal contact

To better understand the potential for mucosal transmission reported for the current Clade I outbreak, we simulated a mucosal contact transmission event. We designed the experiment to compare the transmission efficiency of rectal versus nasal mucosal contact. For this, we used a polyester-tipped sampling swab to transfer virus-containing mucus and debris from the mucosal surface of a donor to the mucosa of a naïve sentinel animal during peak shedding (Figure 5A). Donor animals were rectally or nasally inoculated (*n=4* transmission pairs per route). On day 11 post exposure, a swab sample was taken from the donor’s inoculation site (rectum or nose) and inserted into the respective mucosal site of a naïve sentinel. Sentinels received either a rectal swab from a rectally infected donor, or a nasal swab from a nasally infected donor. At the end of the simulated transmission event, transfer-swab samples were diluted in media and subjected to virus titration to estimate the residual viral load on the swabs and assess the efficiency of simulated transmission. Titration of the swabs after the transmission revealed a titer of 3-5.5×10^6^ PFU/ml for rectal and 1.25×10^5^-2.75×10^6^ PFU/ml for nasal transfer swabs (Supplemental Figure 5).

**Figure 5:**
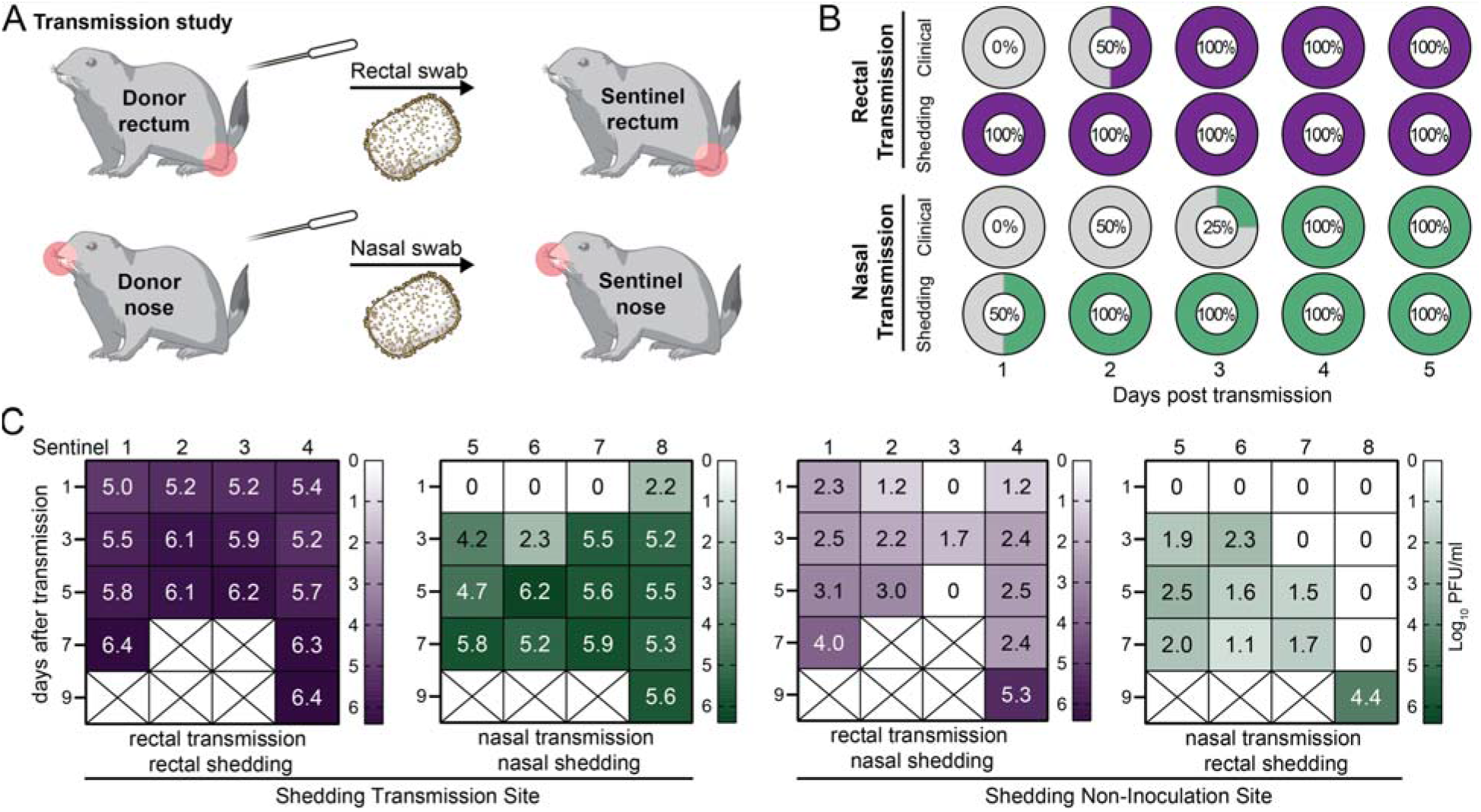
Rapid onset of virus shedding after MPXV transmission. (A) Simulated mucosal contact transmission. Donor animals were nasally or rectally infected. 11 days post inoculation, virus-containing mucus was taken from the inoculation site and transferred to the respective orifice of a naïve sentinel. From the rectally inoculated donor, a rectal swab was used for the transfer and from the nasally inouclated donor, a nasal swab was used. (B) Temporal connection between onset of clinical signs and infectious virus shedding in sentinel animals (*n=4*) after rectal or nasal transmission. Percentage of animals displaying clinical signs during exam, including localized inflammation (swelling or redness), or generalized lesions. Percentage of animals with at least one positive swab sample (oropharyngeal, nasal, urogenital, or rectal) with infectious virus. Animals that reached humane endpoint criteria after productive virus shedding were counted as positive for the remaining days. (C-F) Infectious virus shedding in sentinel animals after transmission. Virus titer in Log_10_ PFU/ml is displayed for different days after transmission. Panel C and D show virus shedding at exposed mucosa. E and F show shedding at non-transmission sites (rectal shedding after nasal transmission and nasal shedding after rectal transmission). A cross indicates the animal had reached humane endpoint criteria and was removed from the study.

Overall transmission efficiency was 100%, with all sentinels reaching endpoint criteria within 9 days after severe systemic infection (Supplemental Figure 6). After the rectal transmission event, all sentinel animal started shedding infectious virus (1×10^5^-2.25×10^5^ PFU/ml) at contact mucosa already on day 1. Of the four nasal transmission sentinels, only one animal shed infectious virus (1.75×10^2^ PFU/ml) on day 1 at their contact mucosa. When testing for virus shedding at non-transmission sites, virus loads were significantly lower compared to the inoculation site. 3 out of the 4 sentinels after rectal transmission had infectious virus in nasal swabs on day 1, with 2×10^2^ PFU/ml as highest value. In contrast, none of the sentinels after nasal contact showed shedding in rectal swabs on day 1. Compared to the rectal transmission group, viral shedding from non-transmission sites was significantly lower in the nasal transmission group.

Although transmission was observed via both routes, rectal transmission was more efficient than nasal transmission, as evidenced by higher viral loads at both the transmission site and distal (non-transmission) mucosal sites.

### Infectious virus shedding occurs before first clinical signs

First clinical signs after rectal transmission included redness, and swelling of the anus, which was first detected on day 3 after transmission in 2 out of 4 sentinels. All sentinels of the rectal groups showed clinical signs on day 5. Histologically, acute, severe anal and rectal ulceration, necrosis, and inflammation were observed (Figure 6). After nasal transmission, nasal discharge was first observed in 1 out of 4 animals after 5 days and in all after 7 days. Nasally exposed sentinels developed predominantly respiratory clinical signs, including dyspnea due to obstructed nasal turbinates. The nasal turbinate tissue of these animals showed an acute rhinitis (Figure 6). All These first clinical signs were detected 2-4 days after the onset of infectious virus shedding (Figure 5B), indicating pre-symptomatic shedding for rectal and nasal transmission.

**Figure 6:**
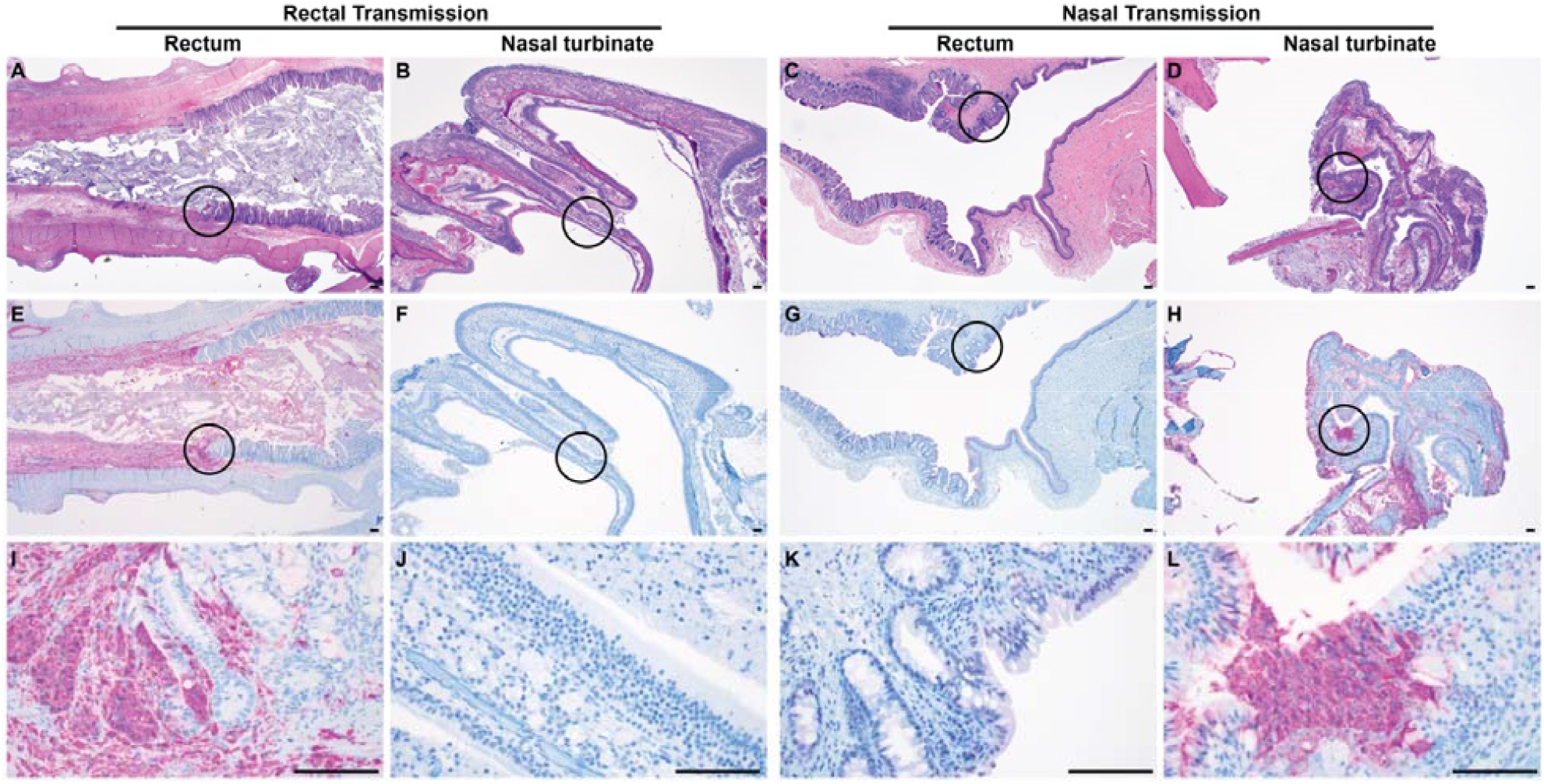
Histologic lesions of sentinel animals after rectal or nasal transmission demonstrate inflammation and necrosis. The first column of panels (A,E,I) shows rectal mucosa after rectal transmission, the second column (B,F,J) shows nasal mucosa after rectal transmission. Rectal mucosa displayed ulceration with subjacent submucosal inflammation, fibroplasia and edema of the submucosa extending to the muscularis. Glandular epithelial cells, submucosal reactive fibroblasts, endothelial cells and inflammatory cells show antigen presence in IHC. Nasal mucosa does not display any pathological changes and does not present with MPXV antigen positivity in IHC. Panel column three displays rectal mucosa after nasal transmission (C,G,K) and the last column displays nasal mucosa after nasal transmission (D,H,L). After nasal transmission, no pathological changes or MPXV antigen presence were detected in the mucosa of the anorectal junction. Nasal turbinates presented with inflammation of a portion of the olfactory epithelial submucosa and corresponding foci of epithelial cells with MPXV antigen. Panels A-D are hematoxylin and eosin stained (40x magnification), E-H (40x magnification) and I-L (400x magnification) are immunohistochemistry (IHC) images using a polyclonal IgG vaccinia virus antibody. Scale bars indicate 100µm.

## Discussion

Epidemiological data indicate close physical contact, including sexual interactions, as the key driver of sustained human-to-human transmission during the 2022 and the ongoing mpox outbreaks^27-29^. However, knowledge gaps remain regarding the efficient spread of the currently circulating variants. Here, we investigated MPXV clade Ia pathogenicity, disease progression, shedding kinetics, and transmission efficiency following different mucosal routes of infection utilizing the black-tailed prairie dog (*Cynomys ludovicianus*) model. Former studies have demonstrated the transmissibility of MPXV clade IIa between prairie dogs via fomites, respiratory/aerosolized fomites, or co-housing^24,25^. Here, we demonstrated that MPXV causes highly productive infections after anogenital mucosal exposure and can be efficiently transmitted via mucosal contact.

Animals exposed at their anogenital (penile, vaginal, or rectal) mucosa displayed a more severe disease progression compared to the intranasally challenged group. Signs of more pronounced clinical disease included increased virus spread and replication in systemic organs and stronger immune cell activation. These results coincide with findings in multimammate rats (*Mastomys natalensis*), where rectal or vaginal challenge with MPXV clade IIb resulted in increased virus shedding compared to intraperitoneal, transdermal, or aerosol exposure^30^. The higher susceptibility of anogenital mucosa might be associated with a different composition of the epithelium of the respective mucosa, including tissue resident immune cells^31-33^.

As shown for other virus infections like HIV or HSV, inflicting damage to the mucosal barrier can facilitate successful establishment of infections^34-36^. Mucosal barriers including mucus and cilia acts as a defense mechanism against several viral infections^37,38^. Additionally, the inflicted microabrasions strip away upper layers of the stratum corneum, which function as a mechanical barrier against pathogens. Similarly, for other viruses that enter via mucosal surfaces, the disruption of the tight-junction barrier can facilitate virus entry by exposing vulnerable basolateral membranes with extracellular and transmembrane proteins that can act as virus receptors^34^. Sexual contact can disrupt the integrity of mucosal barriers and cause microabrations, which can facilitate the virus entry and can cause a productive infection. Whereas it is likely that these mechanisms play a role in the efficient transmission of mpox, the exact impact of the inflicted micro-damages on the epithelial barrier needs further investigation.

Yet, this study has potential limitations. Mucus removal and inflicting microabrasions has been performed on the rectum, vagina, penis/preputium, and the nostrils before inoculation. The fact that the mucosa of the nostrils and not the mucosa in the nasal cavity has been treated prior to inoculation, can have contributed to the milder disease we detected after nasal exposure. Moreover, it must be taken into consideration that these results were generated in an animal model and translation to the current outbreak situation must be done carefully.

Here, we describe a rapid disease progression and more severe clinical disease in sentinel animals after mucosal transmission compared to experimentally inoculated animals. The exact exposure dose of the sentinel animals during the transmission events is challenging to quantify. Still, the titration data of the transfer swabs indicate high-titer exposure and accentuates the virus-richness of mucus at the infection site, which can facilitate efficient virus infection from one mucosal site to the next. However, additional factors may also play a role, such as virions released from the mucosal surfaces of the donor may be more infectious than *in vitro* grown virus. Naturally released virions (extracellular enveloped virions) might benefit from their additional outer membrane including different glycosylation pattern, which is missing on infectious virions artificially extracted from cell culture (intracellular mature virions)^39,40^. This outer membrane, which is important for efficient intra-host cell-to-cell spread, might affect the transmission after mucosal contact. Additionally, if and how the exposure to cellular debris, chemokines and cytokines, and mucus from the donor animal’s mucosa alters the host’s immune response and facilitate or inhibit virus entry demands further investigations. Similar findings have been reported in a former prairie dog transmission study where contact sentinels developed a more severe disease progression compared to donors^24^.

The preference of mpox for mucosal contact transmission has so far predominantly been reported for clade Ib and IIb outbreaks, especially in the context of sexual interactions^27-29^. However, this study demonstrates the ability of clade Ia to have the same propensity to be transmitted by mucosal contact. This indicates the mucosal preference seems to be present in all mpox clades, with a similarly high potential of clade Ia for effective spread after close contact. Virulence likely varies between mpox clades and even individual strains, with milder variants causing only localized lesions. But the general transmission mode might depend on anthropogenic behavior and is less clade-specific feature. Additionally, our results show a systemic spread following mucosal infection, but with high virus shedding and severe inflammatory processes occurring at the infection site.

A key finding of this study is the observation of early, pre-symptomatic and high-titer virus shedding after rectal transmission. The rapid, pre-symptomatic shedding of infectious virus was only observed at the site of initial virus entry, suggesting that mpox transmission from infected mucosa can occur before the systemic dissemination or the occurrence of the classic skin lesions. Therefore, shedding and transmission of infectious virus before patients develop clinical symptoms could be an important driver for transmission in the ongoing outbreak. The detection of ABOBEC3 mutations in clade Ia strains from the Democratic Republic of the Congo and the Republic of the Congo indicate previously unrecognized human-to-human transmission in central Africa^41-43^, demonstrating the potential for pre- or asymptomatic transmission chains. Further investigation into the role of pre-symptomatic mpox transmission in the current outbreak could provide a better understanding of the rapid mpox spread throughout Africa and beyond.

In our animal model, generalized skin lesions appear relatively late during the disease, several days after the onset of virus shedding or localized inflammation. At that time point, the infection has progressed to a systemic dissemination with all organs being affected. This coincides with data about disease progression in humans, where macules (the first stage of skin lesions) appear after an incubation period and a prodromal disease phase that manifests by fever, malaise, and lymphadenopathy ^44,45^. The relatively late appearance of disseminated skin lesions, and given that viremia at the timepoint of lesion onset is an important biomarker for disease severity^46^, indicate the need to ideally initiate antiviral treatment before the occurrence of generalized skin lesions. This may explain the poor performance of the antiviral Tecovirimat during the PALM007 trial^47^. Additionally, patients vaccinated with pre-exposure JYNNEOS were protected against severe systemic disease but still displayed localized infections, predominately in their genital area after contracting an mpox clade IIb infection^48^. Based on the significant role of genital and rectal mpox infections and the highly efficient anogenital transmission demonstrated in this study, we advocate for a stronger focus on anogenital infections during preclinical and clinical evaluations of countermeasures.

Overall, this study proves the increased shedding kinetics and pathogenicity following anogenital infections and simulated mucosal contact transmission in an *in vivo* model. Our results demonstrate the potential of mpox to cause rapid and high-titer virus shedding after anogenital transmission events. These results have important implications for understanding human transmission chains and build a foundation for future research to further improve mpox outbreak responses.

## Data and materials availability

All data supporting the findings of this study have been deposited on Figshare and will be made publicly available upon acceptance. A DOI will be provided in the final published version.

## Acknowledgements

We thank the Poxvirus and Rabies Branch at the Centers for Disease Control and Prevention (CDC), Georgia, USA, for helpful discussion; BEI, NIAID resources for supplying mpox isolates; the Office of the Chief, RML, NIAID, NIH for their support within the high containment facility; the animal care staff and veterinarians of the Rocky Mountain Veterinary Branch, NIAID, NIH for their assistance during the study; and the Visual and Medical Arts Section, RML, NIAID, NIH for their help in preparing the graphical illustration.

## References

1. Rimoin AW, Mulembakani PM, Johnston SC, et al. Major increase in human monkeypox incidence 30 years after smallpox vaccination campaigns cease in the Democratic Republic of Congo. Proc Natl Acad Sci U S A 2010;107(37):16262–7. (In eng). DOI: 10.1073/pnas.1005769107.

2. Ndembi N, Folayan MO, Ngongo N, et al. Mpox outbreaks in Africa constitute a public health emergency of continental security. Lancet Glob Health 2024;12(10):e1577–e1579. (In eng). DOI: 10.1016/s2214-109x(24)00363-2.

3. Likos AM, Sammons SA, Olson VA, et al. A tale of two clades: monkeypox viruses. J Gen Virol 2005;86(Pt 10):2661–2672. (In eng). DOI: 10.1099/vir.0.81215-0.

4. Chen N, Li G, Liszewski MK, et al. Virulence differences between monkeypox virus isolates from West Africa and the Congo basin. Virology 2005;340(1):46–63. (In eng). DOI: 10.1016/j.virol.2005.05.030.

5. Moss B. Understanding the biology of monkeypox virus to prevent future outbreaks. Nature Microbiology 2024;9(6):1408–1416. DOI: 10.1038/s41564-024-01690-1.

6. Adler H, Gould S, Hine P, et al. Clinical features and management of human monkeypox: a retrospective observational study in the UK. Lancet Infect Dis 2022;22(8):1153–1162. (In eng). DOI: 10.1016/s1473-3099(22)00228-6.

7. Gigante CM, Korber B, Seabolt MH, et al. Multiple lineages of monkeypox virus detected in the United States, 2021-2022. Science 2022;378(6619):560–565. (In eng). DOI: 10.1126/science.add4153.

8. Bangwen E, Diavita R, De Vos E, et al. Suspected and confirmed mpox cases in DR Congo: a retrospective analysis of national epidemiological and laboratory surveillance data, 2010–23. The Lancet 2025;405(10476):408–419. DOI: 10.1016/S0140-6736(24)02669-2.

9. Ndembi N, Folayan MO, Komakech A, et al. Evolving Epidemiology of Mpox in Africa in 2024. N Engl J Med 2025 (In eng). DOI: 10.1056/NEJMoa2411368.

10. Organization WH. WHO Director-General declares mpox outbreak a public health emergency of international concern. (https://www.who.int/news/item/14-08-2024-who-director-general-declares-mpox-outbreak-a-public-health-emergency-of-international-concern).

11. Wawina-Bokalanga T, Akil-Bandali P, Kinganda-Lusamaki E, et al. Co-circulation of monkeypox virus subcladesl1Ia and Ib in Kinshasa Province, Democratic Republic of the Congo, July to August 2024. Euro Surveill 2024;29(38) (In eng). DOI: 10.2807/1560-7917.Es.2024.29.38.2400592.

12. CDC. Mpox in the United States and Around the World: Current Situation. 2/11/2025 (https://www.cdc.gov/mpox/situation-summary/index.html).

13. O’Toole Á, Neher RA, Ndodo N, et al. APOBEC3 deaminase editing in mpox virus as evidence for sustained human transmission since at least 2016. Science 2023;382(6670):595–600. (In eng). DOI: 10.1126/science.adg8116.

14. Vakaniaki EH, Kacita C, Kinganda-Lusamaki E, et al. Sustained human outbreak of a new MPXV clade I lineage in eastern Democratic Republic of the Congo. Nature Medicine 2024;30(10):2791–2795. DOI: 10.1038/s41591-024-03130-3.

15. Brosius I, Vakaniaki EH, Mukari G, et al. Epidemiological and clinical features of mpox during the clade Ib outbreak in South Kivu, Democratic Republic of the Congo: a prospective cohort study. The Lancet. DOI: 10.1016/S0140-6736(25)00047-9.

16. Nations U. Eastern DR Congo crisis increasing risk of mpox transmission, WHO chief warns. 2/3/2025 (https://news.un.org/en/story/2025/02/1159701).

17. Weiner ZP, Salzer JS, LeMasters E, et al. Characterization of Monkeypox virus dissemination in the black-tailed prairie dog (Cynomys ludovicianus) through in vivo bioluminescent imaging. PLoS One 2019;14(9):e0222612. (In eng). DOI: 10.1371/journal.pone.0222612.

18. Hutson CL, Olson VA, Carroll DS, et al. A prairie dog animal model of systemic orthopoxvirus disease using West African and Congo Basin strains of monkeypox virus. J Gen Virol 2009;90(Pt 2):323–333. (In eng). DOI: 10.1099/vir.0.005108-0.

19. Americo JL, Earl PL, Moss B. Virulence differences of mpox (monkeypox) virus clades I, IIa, and IIb.1 in a small animal model. Proceedings of the National Academy of Sciences 2023;120(8):e2220415120. DOI: 10.1073/pnas.2220415120.

20. Earl PL, Americo JL, Moss B. Insufficient Innate Immunity Contributes to the Susceptibility of the Castaneous Mouse to Orthopoxvirus Infection. J Virol 2017;91(19) (In eng). DOI: 10.1128/jvi.01042-17.

21. Hutson CL, Kondas AV, Mauldin MR, et al. Pharmacokinetics and Efficacy of a Potential Smallpox Therapeutic, Brincidofovir, in a Lethal Monkeypox Virus Animal Model. mSphere 2021;6(1) (In eng). DOI: 10.1128/mSphere.00927-20.

22. Keckler MS, Salzer JS, Patel N, et al. IMVAMUNE® and ACAM2000® Provide Different Protection against Disease When Administered Postexposure in an Intranasal Monkeypox Challenge Prairie Dog Model. Vaccines.

23. Smith SK, Self J, Weiss S, et al. Effective antiviral treatment of systemic orthopoxvirus disease: ST-246 treatment of prairie dogs infected with monkeypox virus. J Virol 2011;85(17):9176–87. (In eng). DOI: 10.1128/jvi.02173-10.

24. Hutson CL, Carroll DS, Gallardo-Romero N, et al. Monkeypox disease transmission in an experimental setting: prairie dog animal model. PLoS One 2011;6(12):e28295. (In eng). DOI: 10.1371/journal.pone.0028295.

25. Guarner J, Johnson BJ, Paddock CD, et al. Monkeypox transmission and pathogenesis in prairie dogs. Emerg Infect Dis 2004;10(3):426–31. (In eng). DOI: 10.3201/eid1003.030878.

26. Li Y, Zhao H, Wilkins K, Hughes C, Damon IK. Real-time PCR assays for the specific detection of monkeypox virus West African and Congo Basin strain DNA. J Virol Methods 2010;169(1):223–7. (In eng). DOI: 10.1016/j.jviromet.2010.07.012.

27. Kibungu E, Vakaniaki E, Kinganda-Lusamaki E, et al. Clade I–Associated Mpox Cases Associated with Sexual Contact, the Democratic Republic of the Congo. Emerging Infectious Disease journal 2024;30(1). DOI: 10.3201/eid3001.231164.

28. Katoto PD, Muttamba W, Bahizire E, et al. Shifting transmission patterns of human mpox in South Kivu, DR Congo. Lancet Infect Dis 2024;24(6):e354–e355. (In eng). DOI: 10.1016/s1473-3099(24)00287-1.

29. de Vries HJ, Götz HM, Bruisten S, et al. Mpox outbreak among men who have sex with men in Amsterdam and Rotterdam, the Netherlands: no evidence for undetected transmission prior to May 2022, a retrospective study. Eurosurveillance 2023;28(17):2200869. DOI: doi:10.2807/1560-7917.ES.2023.28.17.2200869.

30. Port JR, Riopelle JC, Smith SG, et al. Infection with mpox virus via the genital mucosae increases shedding and transmission in the multimammate rat (Mastomys natalensis). Nature Microbiology 2024;9(5):1231–1243. DOI: 10.1038/s41564-024-01666-1.

31. Gonzalez SM, Aguilar-Jimenez W, Su R-C, Rugeles MT. Mucosa: Key Interactions Determining Sexual Transmission of the HIV Infection. Frontiers in Immunology 2019;Volume 10 - 2019 (Review) (In English). DOI: 10.3389/fimmu.2019.00144.

32. Bomsel M, Alfsen A. Entry of viruses through the epithelial barrier: pathogenic trickery. Nat Rev Mol Cell Biol 2003;4(1):57–68. (In eng). DOI: 10.1038/nrm1005.

33. Yüzen D, Arck PC, Thiele K. Tissue-resident immunity in the female and male reproductive tract. Semin Immunopathol 2022;44(6):785–799. (In eng). DOI: 10.1007/s00281-022-00934-8.

34. Yoon M, Spear Patricia G. Disruption of Adherens Junctions Liberates Nectin-1 To Serve as Receptor for Herpes Simplex Virus and Pseudorabies Virus Entry. Journal of Virology 2002;76(14):7203–7208. DOI: 10.1128/jvi.76.14.7203-7208.2002.

35. Rana H, Truong NR, Sirimanne DR, Cunningham AL. Breaching the Barrier: Investigating Initial Herpes Simplex Viral Infection and Spread in Human Skin and Mucosa. Viruses.

36. Burgener A, McGowan I, Klatt NR. HIV and mucosal barrier interactions: consequences for transmission and pathogenesis. Curr Opin Immunol 2015;36:22–30. (In eng). DOI: 10.1016/j.coi.2015.06.004.

37. Lieleg O, Lieleg C, Bloom J, Buck CB, Ribbeck K. Mucin biopolymers as broad-spectrum antiviral agents. Biomacromolecules 2012;13(6):1724–32. (In eng). DOI: 10.1021/bm3001292.

38. Habte HH, Mall AS, de Beer C, Lotz ZE, Kahn D. The role of crude human saliva and purified salivary MUC5B and MUC7 mucins in the inhibition of Human Immunodeficiency Virus type 1 in an inhibition assay. Virology Journal 2006;3(1):99. DOI: 10.1186/1743-422X-3-99.

39. Moss B. Poxvirus cell entry: how many proteins does it take? Viruses 2012;4(5):688–707. (In eng). DOI: 10.3390/v4050688.

40. Condit RC. Surf and turf: mechanism of enhanced virus spread during poxvirus infection. Viruses 2010;2(5):1050–1054. (In eng). DOI: 10.3390/v2051050.

41. Wawina-Bokalanga T, Merritt S, Kinganda-Lusamaki E, et al. Epidemiology and Phylogenomic Characterization of Distinct 2023 and 2024 Mpox outbreaks in Kinshasa, Democratic Republic of the Congo - Evidence for increasingly sustained human-to-human transmission of subclade Ia. medRxiv 2024:2024.11.15.24317404. DOI: 10.1101/2024.11.15.24317404.

42. Otieno JR, Ruis C, Onoja AB, et al. Global genomic surveillance of monkeypox virus. Nat Med 2025;31(1):342–350. (In eng). DOI: 10.1038/s41591-024-03370-3.

43. Yinda CK, Koukouikila-Koussounda F, Mayengue PI, et al. Genetic sequencing analysis of monkeypox virus clade I in Republic of the Congo: a cross-sectional, descriptive study. The Lancet 2024;404(10465):1815–1822. DOI: 10.1016/S0140-6736(24)02188-3.

44. Di Giulio DB, Eckburg PB. Human monkeypox: an emerging zoonosis. Lancet Infect Dis 2004;4(1):15–25. (In eng). DOI: 10.1016/s1473-3099(03)00856-9.

45. Breman JG, Kalisa R, Steniowski MV, Zanotto E, Gromyko AI, Arita I. Human monkeypox, 1970-79. Bull World Health Organ 1980;58(2):165–82. (In eng).

46. Nishiyama T, Miura F, Jeong YD, et al. Modeling lesion transition dynamics to clinically characterize patients with clade I mpox in the Democratic Republic of the Congo. Sci Transl Med 2025;17(805):eads4773. (In eng). DOI: 10.1126/scitranslmed.ads4773.

47. Tecovirimat for Clade I MPXV Infection in the Democratic Republic of Congo. New England Journal of Medicine 2025;392(15):1484–1496. DOI: doi:10.1056/NEJMoa2412439.

48. Granskog L, Saadeh K, Lorenz K, et al. Effect of JYNNEOS vaccination on mpox clinical progression: a case-control study. Lancet Infect Dis 2025 (In eng). DOI: 10.1016/s1473-3099(25)00180-x.

