## Supplemental data for "Highly efficient anogenital transmission of clade Ia mpox virus associated with increased shedding"

**Supplemental Figures:**

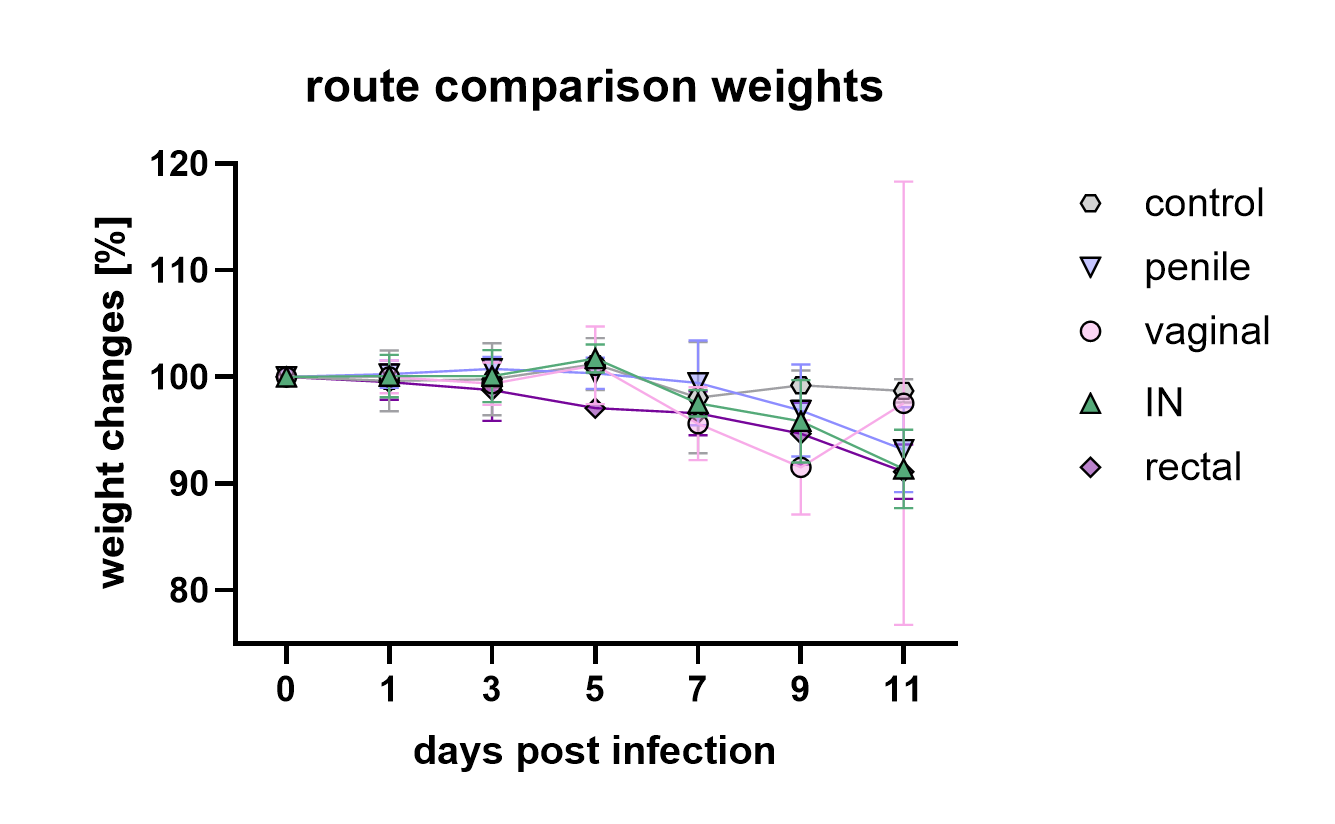

**Supplemental Figure 1. Body weight changes in dependence of inoculation route after MPXV exposure (*n=4*)**. Symbols show means and error bars indicate standard deviation.

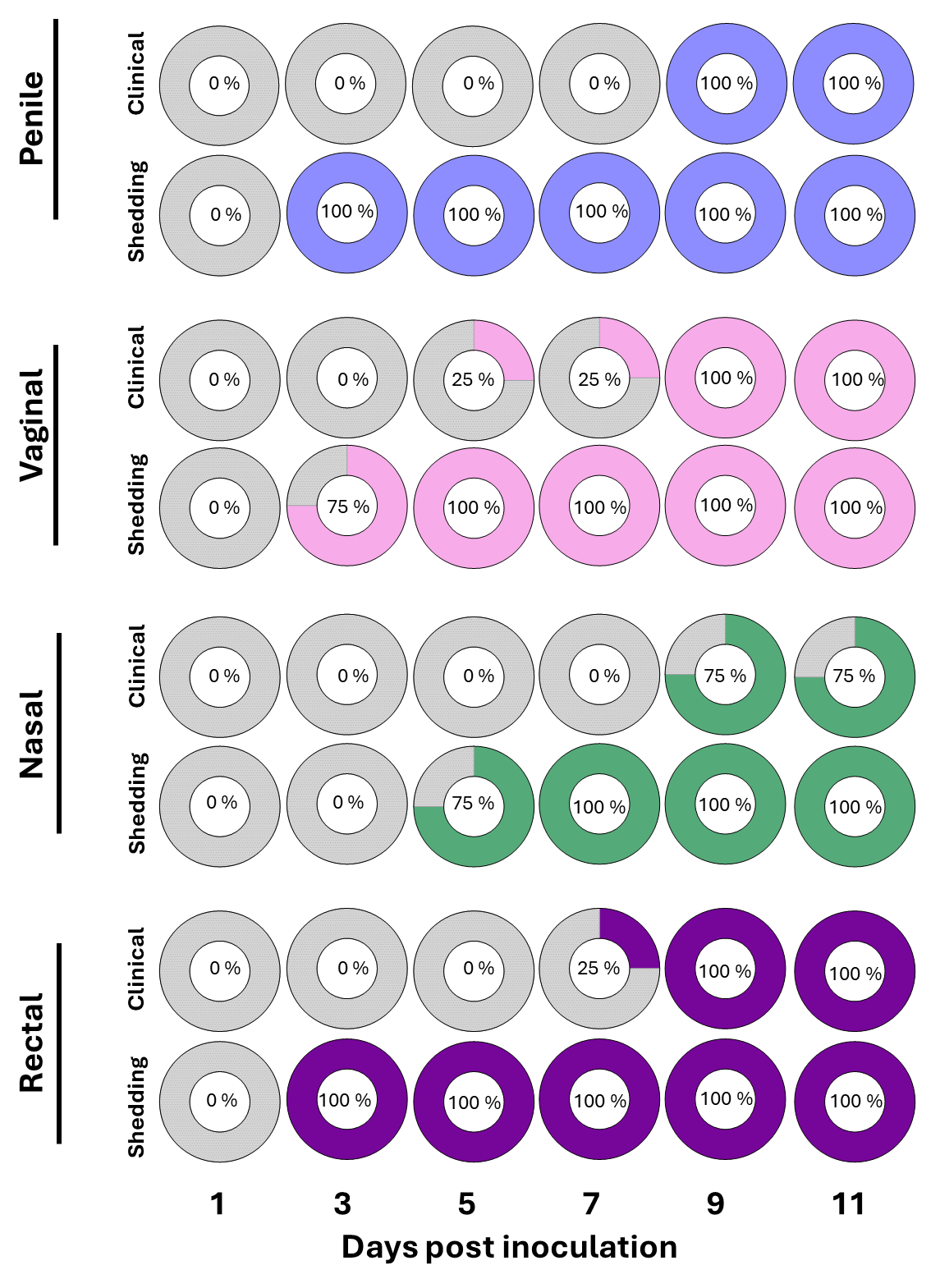

**Supplemental Figure 2: Temporal connection between onset of clinical signs and infectious virus shedding after different exposure routes**. Animals (*n=4*) after penile, vaginal, nasal, or rectal inoculation. Percentage of animals displaying clinical signs during exam, including localized inflammation (swelling or redness), or generalized lesions. Percentage of animals with at least one positive swab sample (oropharyngeal, nasal, urogenital, or rectal) with infectious virus.

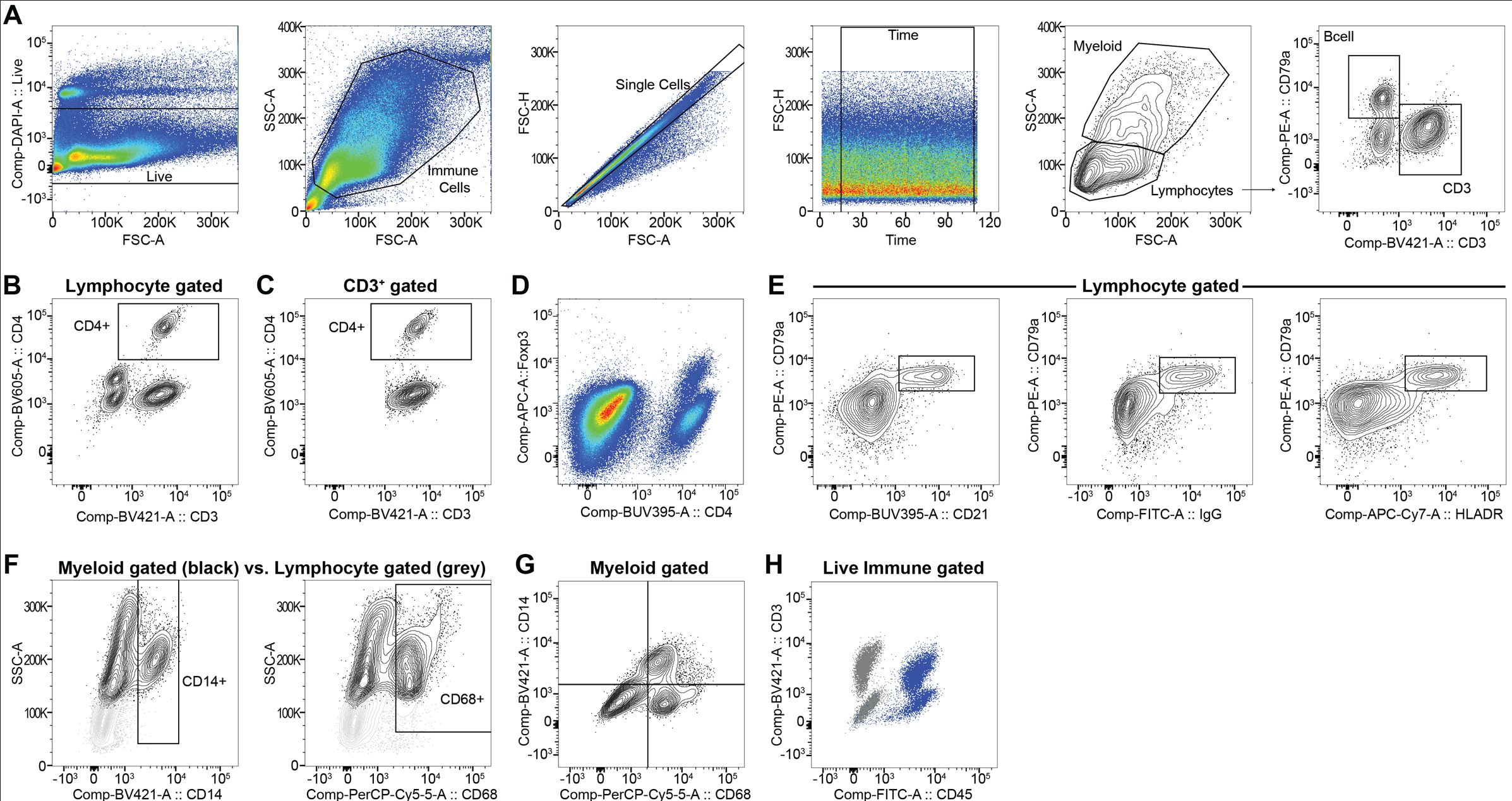

**Supplemental Figure 3. Representative gating strategies for flow analysis of splenocytes from prairie dogs infected by MPXV at 11dpi.** (A) Live splenocytes were first gated with a dead cell exclusion gate using Fixable Blue vs FSC-A. Total immune cells were then gated using SSC-A vs FSC-A, followed by a single cell gate using FSC-H vs FSC-A and then a time gate to exclude possible erratic sample flow. This population was further analyzed by SSC-A vs FSC-A to identify the myeloid and lymphocyte immune cells, which were further analyzed by cross-reactive clones that were identified in prairie dogs (see Supplemental Table 1 for clone information). The lymphocytes were stained using intracellular CD79a and an intracellular epitope for CD3. Lymphocytes were further characterized by (B) surface staining with a cross-reactive surface CD4 antibody, which (C) identified CD4^+^ T cells that dually stained with CD3^+^. (D) CD4^+^ cells co-stained with intracellular Foxp3. Lymphocytes were further analyzed for cross-reactive B cell markers and (E) CD79a^+^ cells stained with cross-reactive clones for CD21, IgG and HLA-DR, which further corroborate the identification of B cells. (F) The SSC-A^hi^ myeloid population (black) stained with cross-reactive clones for CD14 and CD68, compared to lymphocyte gated cells (light gray), as depicted in representative over-layed FACS plots. (G) A representative co-staining plot of CD68 and CD14 for myeloid-gated immune cells. (H) A representative overlay plot of total live immune cells stained with live/dead only (gray) and CD45 and CD3 (blue).

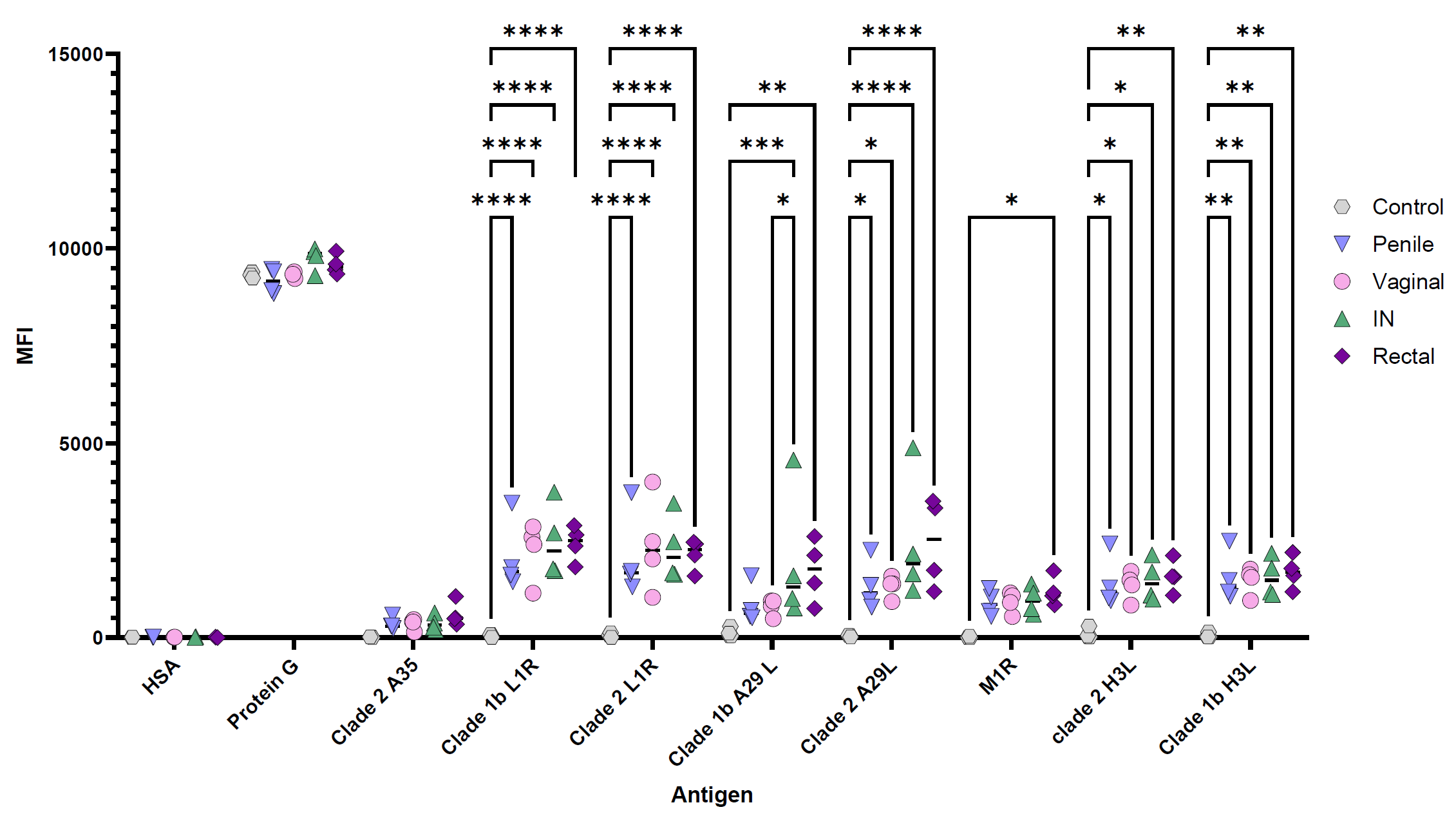
**Supplemental Figure 4: Antibody titers against selected MPXV antigens on day 11 post inoculation.** A semi-quantitative bead based multiplex assay was developed to analyze antibody binding using Luminex MagPlex C microspheres coupled to the indicated recombinant antigen. Protein G beads act as a positive internal control, and beads coupled to human serum albumin act as a negative internal control. Data is shown as median fluorescence intensity (MFI), and route of infection is indicated in figure key. P-values were calculated with a two-way ANOVA and Tukey’s multiple comparisons test. * P < 0.05, ** P < 0.01, *** P < 0.001, **** P < 0.0001

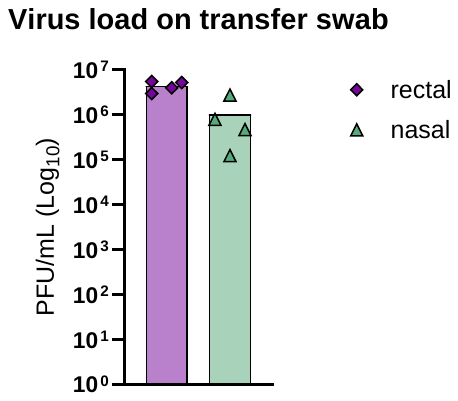

**Supplemental Figure 5. Infectious virus on transfer swabs from donor animals after they have been used to transfer mucus to sentinels**.

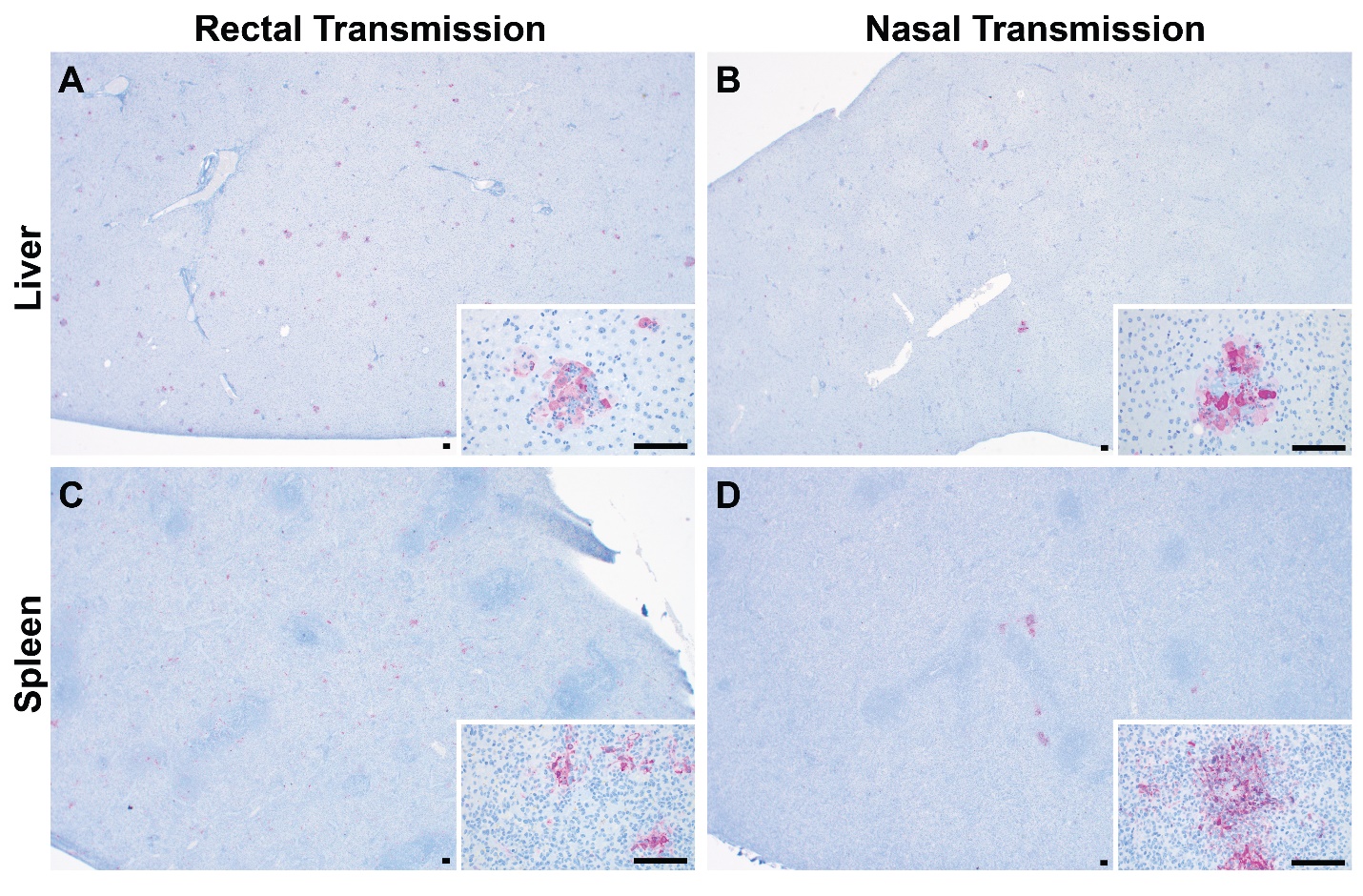

**Supplemental Figure 6: Immunohistochemistry of liver and spleen lesions after systemic virus spread in sentinel animals following rectal or nasal transmission**. All images taken at 20x magnification; all insets at 400x magnification. Scale bars indicate 100µm.

**Supplemental Table 1: Screening process of antibodies for basic immune cell identification and phenotyping**

| **Marker** | **Clone** **yes (gray highlight & bold) *maybe: needs confirmation | **Isotype** | **Company** | **Comments** |
| --- | --- | --- | --- | --- |
| **Leukocyte Common Antigen** |  |  |  |  |
| **CD45** | **** 30-F11** | **Rat IgG2b, κ** | **BD Biosciences** | see Supp. Fig. 3g |
| CD45 | IH-1 | mouse IgG1 | BioRad & Fisher Scientific |  |
| CD45 | HI30 | Mouse IgG1, κ | BioLegend |  |
| CD45 | DO58-1283 | Mouse IgG1, κ | BD Biosciences |  |
| **T cell** |  |  |  |  |
| **CD3** | **** CD3-12** | **Rat / IgG1** | **BioRad** | see Fig. 4c & Supp. Fig.3a,b,c,g |
| CD3 | OKT3 | CAF1 IgG2a, κ | BD Biosciences |  |
| CD3 | 145-2C11 | Armenian Hamster IgG1, κ | BD Biosciences |  |
| TCRβ chain | H57-597 | Armenian Hamster IgG2, ƛ1 | BD Biosciences |  |
| **CD8 T cell** |  |  |  |  |
| CD8 | CT6 | mouse IgG1 | BioRad |  |
| CD8 | 341 | Mouse BALB/c IgG1, κ | BD Biosciences |  |
| CD8 | SK1 | Mouse BALB/c IgG1, κ | BD Biosciences |  |
| CD8 | RPA-T8 | Mouse IgG1, κ | BD Biosciences |  |
| CD8 | 53-6.7 | Rat LOU IgG2a, κ | BD Biosciences |  |
| CD8 | OKT8 | CAF1 IgG2a, κ | BD Biosciences |  |
| CD8 | HIT8a | Mouse IgG1, κ | BD Biosciences |  |
| CD8 | G28 | Mouse IgG2a, κ | BioLegend |  |
| **CD4 T cell** |  |  |  |  |
| **CD4** | **** L200** | **Mouse BALB/c IgG1, κ** | **BD Biosciences** | see Fig.4c & Supp. Fig. 3b,c,d |
| CD4 | CT7 | mouse IgG1 | BioRad |  |
| CD4 | GK1.5 | Rat IgG2b, κ | BioLegend |  |
| CD4 | OKT4 | Mouse IgG2b, κ | BioLegend |  |
| CD4 | RM4-5 | Rat IgG2a, κ | BioLegend |  |
| CD4 | OX-35 | Mouse BALB/c IgG2a, κ | BD Biosciences |  |
| CD4 | W3/25 | Mouse BALB/c IgG2a, κ | BD Biosciences |  |
| CD4 | OX-38 | Mouse BALB/c IgG2a, κ | BD Biosciences |  |
| **Regulatory T cell (Foxp3)** |  |  |  |  |
| **Foxp3** | ** FJK-16s | Rat / IgG2a, kappa | Invitrogen | see Supp. Fig. 3d |
| Foxp3 | * 150D | Mouse IgG1, κ | BioLegend | possible signal but requires further validation |
| Foxp3 | 259D/C7 | Mouse BALB/c IgG1 | BD Biosciences |  |
| Foxp3 | PCH101 | Rat IgG2a, κ | Invitrogen |  |
| **B cell** |  |  |  |  |
| CD19 | 1D3/CD19 | Rat IgG2a, κ | BioLegend |  |
| CD19 | 6D5 | Rat IgG2a | BioLegend |  |
| CD19 | SJ25C1 | Mouse IgG1, κ | BioLegend |  |
| CD19 | HIB19 | Mouse IgG1, κ | BioLegend |  |
| CD20 | 2H7 | Mouse IgG2b, κ | BD Biosciences |  |
| **CD21** | **** B-ly4** | **Mouse IgG1, κ** | **BD Biosciences** | see Supp. Fig. 3e |
| CD24 | M1/69 | Rat IgG2b, κ | BioLegend |  |
| **CD79a** | **** HM47** | **Mouse BALB/c IgG1, κ** | **BD Biosciences** | see Fig 4c & Supp. Fig. 3a,e |
| B220/CD45R | RA3-6B2 | Rat IgG2a, κ | BioLegend |  |
| anti-Guinea Pig IgG (H/L) | * goat polyclonal IgG | goat polyclonal IgG | BioRad | possible signal but requires further validation |
| IgD | 11-26c.2a | Rat IgG2a, κ | BD Biosciences |  |
| IgM | II/41 | Rat IgG2a, κ | BD Biosciences |  |
| **Myeloid Cells** |  |  |  |  |
| CD11b | M1/70 | Rat IgG2b, κ | BioLegend |  |
| CD11b | ICRF44 | Mouse IgG1, κ | BioLegend |  |
| CD11c | HL3 | Armenian Hamster IgG1, λ2 | BD Biosciences |  |
| CD11c | N418 | Armenian Hamster IgG2 | BD Biosciences |  |
| CD11c | 3.9 | Mouse IgG1, κ | BD Biosciences |  |
| CD11b/c | * OX-42 | Mouse IgG2a, κ | BioLegend | possible signal but requires further validation |
| **CD14** | **** Tuk4** | **Mouse IgG2a** | **Invitrogen** | see Supp. Fig. 3f |
| CD14 | M5E2 | Mouse IgG2a, κ | BD Biosciences |  |
| CD14 | rmC5-3 | Rat LOU IgG1, κ | BD Biosciences |  |
| CD14 | Sa14-2 | Rat IgG2a, κ | BD Biosciences |  |
| CD14 | 61D3 | Mouse IgG1, κ | Invitrogen |  |
| **CD68** | **** FA-11** | **Rat IgG2a** | **BioLegend** | see Supp. Fig. 3f |
| CD68 | ED1 | mouse IgG1 | Novus |  |
| F4/80 | BM8 | Rat IgG2a | BioLegend |  |
| Ly6G/Ly6C (GR1) | RB6-8C5 | Rat IgG2b | BioLegend |  |
| Ly6G | IA8 | Rat / IgG2a, κ | BD Biosciences |  |
| SIRP⍺ | P84 | Rat IgG1, κ | BioLegend |  |
| **Natural Killer Cells** |  |  |  |  |
| NK1.1 | PK136 | Mouse IgG2a, κ | BioLegend |  |
| CD56 | HCD56 | Mouse IgG1, κ | BioLegend |  |
| CD93 | AA4.1 | Rat IgG2b, κ | BioLegend |  |
| CD94 | 18d3 | Rat IgG2a, κ | BioLegend |  |
| **Major Histocompatibility Complex** |  |  |  |  |
| **HLA-DR** | **** LN3** | **Mouse IgG2b, κ** | **BioLegend** | see Fig 4H & Supp. Fig. 3e |
| HLA-DR | L243 | Mouse IgG2a, κ | BioLegend |  |
| MHCII (I-Ek) | 14-4-4s | Mouse IgG2a, κ | BD Biosciences |  |
| MHCII (I-A/I-E) | M5/114.15.2 | Rat IgG2b, κ | BD Biosciences |  |
| MHCII (I-A/I-E) | 2G9 | DA/HA IgG2a, κ | BD Biosciences |  |
| **Cell Phenotyping** |  |  |  |  |
| CD11a | 2D7 | Rat IgG2a, κ | BioLegend |  |
| CD25 | OX-39 | Mouse IgG1, κ | BioLegend |  |
| CD25 | PC61 | Rat IgG1, κ | BioLegend |  |
| CD27 | * LG.7F9 | Armenian hamster IgG | Invitrogen | possible signal but requires further validation |
| CD27 | * LG.3A10 | Armenian Hamster IgG | BioLegend | possible signal but requires further validation |
| CD27 | * M-T271 | Mouse IgG1, κ | BioLegend | possible signal but requires further validation |
| CD27 | * O323 | Mouse IgG1, κ | BioLegend | possible signal but requires further validation |
| CD28 | CD28.2 | Mouse IgG1, κ | BioLegend |  |
| CD28 | 37.51 | Syrian hamster IgG | BioLegend |  |
| CD40 | * 3/23 | Rat IgG2a, κ | BioLegend | possible signal but requires further validation |
| CD40 | * HB14 | Mouse IgG1, κ | BioLegend | possible signal but requires further validation |
| CD43 | 1B11 | Rat IgG2a, κ | BioLegend |  |
| CD44 | IM7 | Rat IgG2a, κ | BioLegend |  |
| CD45RA | 5H9 | Mouse IgG1, κ | BD Biosciences |  |
| CD45RB | * 16A | Rat IgG2a, κ | BD Biosciences | possible signal but requires further validation |
| CD62L | MEL-14 | Rat IgG2a, κ | BioLegend |  |
| CD62L | OX-85 | Mouse IgG1, κ | BioLegend |  |
| CD69 | H1.2F3 | Armenian Hamster IgG | BioLegend |  |
| CD69 | FN50 | Mouse IgG1, κ | BioLegend |  |
| CD154 (CD40L) | MR1 | Armenian hamster IgG | BioLegend |  |
| CD154 (CD40L) | 24-31 | Mouse IgG1, κ | BioLegend |  |
| CD95 (Fas) | DX2 | Mouse IgG1, κ | BioLegend |  |
| CD134 (OX-40) | OX-86 | Rat IgG1, κ | BioLegend |  |
| CD279 (PD-1) | EH12.2H7 | Mouse IgG1, κ | BioLegend |  |
| CD279 (PD-1) | J43 | Armenian Hamster IgG2, κ | BD Biosciences |  |
| CD304 (neuropilin-1) | 3E12 | Rat IgG2a, κ | BioLegend |  |
| KLRG1 | 2F1/KLRG1 | Syrian hamster IgG | BioLegend |  |
| TIGIT (Vstm3) | 1G9 | Mouse IgG1, κ | BD Biosciences |  |
| **Transcription Factors** |  |  |  |  |
| Tbet | * 4B10 | Mouse IgG1, κ | BD Biosciences | possible signal but requires further validation |
| EOMES | Dan11mag | Rat IgG2a, κ | Invitrogen |  |
| Helios | * 22F6 | Armenian Hamster IgG | BD Biosciences | possible signal but requires further validation |
| **Proliferation** |  |  |  |  |
| **Ki-67** | **** B56** | **Mouse IgG1, κ** | **BD Biosciences** |  |
| Ki-67 | * SolA15 | Rat IgG2a, κ | Invitrogen | possible signal but requires further validation |
| Ki-67 | 16A8 | Rat IgG2a, κ | BioLegend |  |
| **Cytokines** |  |  |  |  |
| IL-2 | JES6-5H4 | Rat IgG2b, κ | BioLegend |  |
| IL-2 | MQ1-17H12 | Rat IgG2a, κ | Invitrogen |  |
| IL-10 | JES5-16E3 | Rat IgG2b, κ | BioLegend |  |
| IFNg | XMG1.2 | Rat IgG1, κ | BioLegend |  |
| IFNg | B27 | Mouse IgG1, κ | BioLegend |  |
| IFNg | 4S.B3 | Mouse IgG1, κ | BioLegend |  |
| IFNg | DB-1 | Mouse IgG1, κ | BioLegend |  |
| anti-Guinea Pig TNFa | * goat polyclonal IgG | polyclonal IgG | BioRad | possible signal but requires further validation |
| TNFa | MAb11 | Mouse IgG1, κ | BioLegend |  |
| TNFa | MP6-XT22 | Rat IgG1, κ | BioLegend |  |
| **Cytotoxic Molecules** |  |  |  |  |
| Granzyme A | * GzA-3G8.5 | Mouse IgG2b, κ | Invitrogen | possible signal but requires further validation |
| Granzyme B | GB11 | Mouse IgG1 | Invitrogen |  |
| Granzyme B | NGZB | Rat IgG2a, κ | Invitrogen |  |
| Granzyme B | QA16A02 | Mouse IgG1, κ | BioLegend |  |
| Perforin | S16009B | Rat IgG2a, κ | BioLegend |  |
| Perforin | B-D48 | Mouse IgG1 | BioLegend |  |
